# CGGBP1 regulates chromatin barrier activity and CTCF occupancy at repeats

**DOI:** 10.1101/593137

**Authors:** Divyesh Patel, Manthan Patel, Umashankar Singh

## Abstract

CGGBP1 is a repeat-binding protein with diverse functions in regulation of gene expression, cytosine methylation, repeat silencing and genomic integrity. CGGBP1 has also been identified as a cooperator factor in histone modifying complexes and as a component of protein complexes that form the enhancer-promoter loops. Here we report that the occupancy of CTCF at repeats and the chromatin barrier function of these repeat sequences depends on CGGBP1. Using ChIP-sequencing for CTC we describe CTCF binding on repetitive DNA. Our results show that CGGBP1 determines the CTCF occupancy preference for repeats over canonical CTCF-motif. By combining CTCF ChIP-sequencing results with ChIP-sequencing for three different kinds of histone modifications (H3K4me3, H3K9me3 and H3K27me3) we uncover insulator-like chromatin barrier activities of the repeat-rich CTCF-binding sites. This work shows that CGGBP1 is a regulator of CTCF occupancy and posits it as a regulator of barrier functions of CTCF-binding sites.

## INTRODUCTION

Human CGGBP1 is a ubiquitously expressed protein with important functions in heat shock stress response, cell growth, proliferation and mitigation of endogenous DNA damage (Agarwal et al., 2016; Singh and Westermark, 2011, 2015; Singh et al., 2011, 2014). CGGBP1 has evolved in the amniotes with >98% conservation in homeotherms (Singh and Westermark, 2015). Yet, the involvement of CGGBP1 in highly conserved cellular processes such as cell cycle, maintenance of genomic integrity and cytosine methylation regulation suggests that CGGBP1 fine-tunes these processes in homeothermic organisms to meet the challenges of their terrestrial habitats. CGGBP1 has no known paralogs in the human genome, its expression in human tissues is ubiquitous (Thul and Lindskog, 2018) and RNAi against CGGBP1 causes G1/S arrest or G2/M arrest (Singh et al., 2011) and heat shock stress response-like gene expression changes with variable effects in different cell lines (Singh et al., 2009, 2011). CGGBP1 acts as a cis-regulator of transcription for tRNA genes, Alu elements (Agarwal et al., 2016), FMR1, CDKN1A, HSF1 (Agarwal et al., 2015; Müller-Hartmann et al., 2000) and cytosine methylation regulatory genes including DNMT1 (Agarwal et al., 2015). However, none of these explain the widespread effects CGGBP1 depletion has on global transcriptome. In cultured normal human fibroblasts CGGBP1 depletion induces gene expression shutdown in a manner that resembles the effects of serum starvation (Agarwal et al., 2016). The mechanisms through which CGGBP1 regulates the genome and the transcriptome remain enigmatic.

Recent reports have shown that CGGBP1 regulates cytosine methylation genome-wide with maximum methylation regulatory effects at Alu and LINE elements in CpG context (Patel et al., 2018). The highly prevalent CHH cytosines however show a CGGBP1-dependent methylation pattern at GC-skew regions, insulators and enhancers (Patel et al., 2018). Interestingly, due to CGGBP1 depletion CHH methylation increases at insulators (characterized as CTCF-binding sites) and decreases at enhancers. Preliminarily, these findings lead to the possibility of a crosstalk between CGGBP1 and CTCF. Interestingly, there is additional evidence to support the possibility that CGGBP1 regulates insulator and enhancer activities. Most prominently, a targeted identification of proteins that structure the enhancer-promoter loops identified CGGBP1 as a partner of CTCF and YY1 (Weintraub et al., 2017). Despite its ubiquitous expression, CGGBP1 was not studied further by Weintraub et al because unlike YY1, CGGBP1 was not identified as a hit in a gene essentiality screen (Wang et al., 2015). Although CTCF binds with high affinity to the CTCF-motif (Cuddapah et al., 2009; Kim et al., 2007; Schmidt et al., 2012), it is clear that CTCF is associated with additional genomic elements that do not contain this motif. Interspersed repeats SVA and Alu-SINEs (Pugacheva et al., 2016) as well as microsatellite repeats serve as binding sites for CTCF (Arnold R; Libby et al., 2008; Plasschaert et al., 2014) as well as CGGBP1. Even CTCF-binding sites that contain the CTCF-binding motifs and are not repeats per se have evolved from Alu-SINEs and related repetitive elements (Wang et al., 2012). Additionally, CGGBP1 and CTCF both exhibit cytosine methylation-sensitive DNA binding at GC-rich sequences (Maurano et al., 2015). Further indications of cooperation between CTCF and CGGBP1 come from findings that both of these proteins interact with a crucial tumor suppressor factor NPM1 (Hein et al., 2015; Yusufzai et al., 2004). CGGBP1 is a proven regulator of ribosomal RNA genes that contain CGG triplet repeats and localize to the nucleoli (Müller-Hartmann et al., 2000). NPM1-CTCF interactions determine the organization of chromatin in the nucleolus such that NPM1 establishes that transcriptionally silent DNA resides in the nucleolar periphery (Yusufzai et al., 2004). NPM1-CTCF interaction is required for insulator activity at many sites in the genome and NPM1 complex with CGGBP1 (Hein et al., 2015). CGGBP1 drives ribosomal RNA synthesis upon growth factor stimulation by silencing Alu repeats (Agarwal et al., 2016). Both CGGBP1 and CTCF are reported to have similar nuclear expression in interphase cells and midbody expression in the mitotic cells (Singh et al., 2011; Zhang et al., 2004). These facts viewed collectively further strengthen the possibility that there is a functional crosstalk between CGGBP1 and epigenomic regulator factors such as CTCF.

In this study we have investigated into the possibilities of interaction and regulation between CTCF and CGGBP1. We show that CGGBP1 and CTCF interact with each other. A systematic analysis of CGGBP1 ChIP-seq data, CTCF ChIP-seq data (ENCODE) and co-immunoprecipitation findings show that the binding sites for the two proteins are in vicinity of each other. By studying CTCF occupancy genome-wide through ChIP-seq under conditions of normal CGGBP1 expression, CGGBP1 knockdown and overexpression, we show that CTCF binds to repeats and canonical CTCF-motifs both. Our analysis reveals that CGGBP1 levels determine the CTCF binding preference between repeats and canonical CTCF-motifs. By combining CTCF ChIP-seq with histone modification ChIP-seq under conditions of normal or knocked-down CGGBP1, we demonstrate that CTCF binding sites regulated by CGGBP1 correspond to chromatin barrier elements with profound effects on H3K9me3 distribution. We report a set of potential insulator sequences that are regulated by a CGGBP1-CTCF axis.

## RESULTS

### CGGBP1 and CTCF colocalize and interact with each other

To test the possibilities of functional crosstalk between CGGBP1 and CTCF, we began by testing the subcellular colocalization of the two proteins. Using immunofluorescence in human fibroblasts, we observed that endogenous CGGBP1 and CTCF both predominantly localized to the nuclei (Figure 1A). Using two different antibodies against CGGBP1 as well as CTCF, similar findings were observed (not shown). By analyzing the normalized immunofluorescence signals for CGGBP1 and CTCF, we observed that the expression of the two proteins were correlated in the nuclei. In the cytoplasm, where the expression of both the proteins were weaker, the signals of CGGBP1 and CTCF were not correlated (Figure 1B). CGGBP1 and CTCF are both present in the midbody (Singh and Westermark, 2011; Zhang et al., 2004) and interestingly, we found their colocalization at midbodies as well (Figure 1C). Co-expression and colocalization of two nuclear proteins is not unexpected. To further know if the colocalizing CGGBP1 and CTCF actually do interact with each other, we performed proximity ligation assays. Again, by using two different antibody pairs for CTCF and CGGBP1 we established that these two proteins form complexes (Figure 1D). Quantification of PLA signals reinforced that CGGBP1 and CTCF complexes are far more prevalent in the nuclei (72.18%) than cytoplasm (27.81%) (Figure 1E).

**Figure 1:**
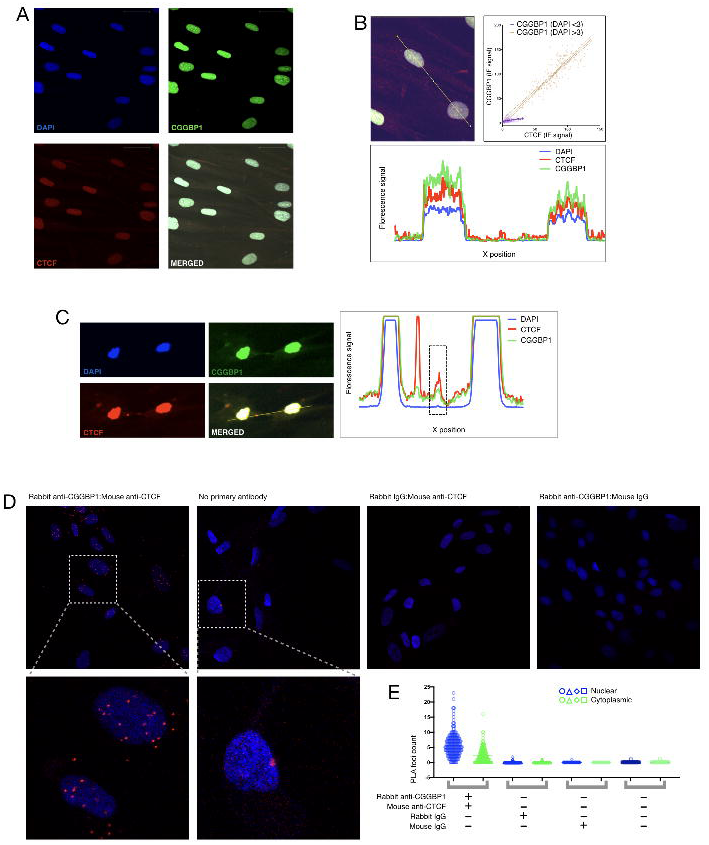
CGGBP1 and CTCF colocalize and interact. **(A)** Human juvenile fibroblasts co-immunostained for CGGBP1 (green) and CTCF (red). Nuclei were counterstained with DAPI (blue). Scale bar = 20 µm. **(B)** Mean fluorescence intensities for CGGBP1 (green) and CTCF (red) were normalised along the line-marked segment and plotted using ImageJ. High correlation was observed between CGGBP1 and CTCF signals in the nuclei (R^2^ for DAPI signals >3 and <3 units are 0.793 and 0.07524 respectively). **(C)** Normalized signals along the line segment drawn through a midbody shows colocalization of CGGBP1 and CTCF. **(D)** PLA (red signal) confirms CTCF-CGGBP1 interaction in situ. Nuclei were stained with DAPI (blue). CGGBP1-CTCF interaction was stronger in the nuclei as than cytoplasm (inset of rabbit anti-CGGBP1:mouse anti-CTCF sample). No significant interaction was observed in IgG and no-primary antibody negative controls (inset of no primary antibody sample). **(E)** Quantification of PLA signals show genuine interactions detected using specific antibodies as highly significantly higher than the negative controls. Also the PLA signal in the nuclei (blue) are significantly higher than the cytoplasm (green).

To further characterize the interactions between endogenously expressed CGGBP1 and CTCF, we performed reciprocal co-immunoprecipitations on whole cell lysates. CGGBP1 expression and function have been well studied in human foreskin fibroblasts. Using CTCF antibodies we could immunoprecipitate CGGBP1 in foreskin fibroblasts. Reciprocally, using CGGBP1 antibodies we could co-precipitate endogenous CTCF and CGGBP1 (Figure 2A). Unlike the rapidly cycling juvenile fibroblasts, the relatively slowly cycling adult dermal fibroblasts did not show a robust pulldown of CTCF and CGGBP1 with each other (not shown). As suggested by IFs and PLA, the interactions were expected more in the nuclear fraction than in the cytoplasmic fraction. CGGBP1 also shows enhanced nuclear presence in growth-stimulated cells as compared to serum-starved cells (Agarwal et al., 2016). We performed co-immunoprecipitations from human dermal fibroblast lysates under conditions of serum starvation and stimulation. In the starved lysates, the reciprocal pulldown of CTCF and CGGBP1 was very weak. In lysates from stimulated cells however, using CGGBP1 antibody we could pull down a major fraction of CTCF (Figure 2B). These findings suggested that the stoichiometry of CTCF-CGGBP1 interactions vary between cell types and also depend on physiological states of the cells. To study functional relevance of CTCF-CGGBP1 interactions we chose to employ knockdown of CGGBP1 using shRNA. Hence, we needed to perform the next set of experiments in human embryonic kidney cells (HEK293T); a cell type that (i) expresses both CTCF and CGGBP1 at detectable levels and (2) in which CGGBP1 knockdown does not induce quiescence. First, we confirmed the interaction between CTCF and CGGBP1 in HEK293T by reciprocal co-immunoprecipitations (Figure 2C). We also subjected the nuclear co-immunoprecipitates to DNAse digestion and characterized the fraction of CGGBP1-CTCF complexes that depend on a DNA bridge for co-immunoprecipitation. We found that there are two kinds of nuclear CGGBP1-CTCF complexes: a smaller fraction that depends on a DNA bridge and another that is direct and does not require a DNA bridge (Figure 2D). Collectively, these findings showed that CGGBP1 and CTCF are present in the nuclei in close proximity, they interact with each other stably and possibly bind to closely located sites on DNA.

**Figure 2:**
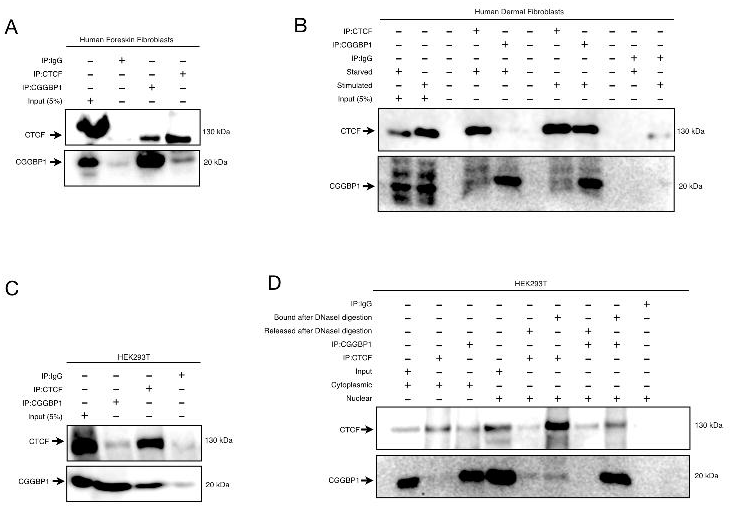
CGGBP1 and CTCF interact variously in different cell types and subcellular fractions. **(A)** Reciprocal co-immunoprecipitations (co-IPs) confirmed interaction between CGGBP1 and CTCF in human foreskin fibroblast. **(B)** Reciprocal co-IPs in human dermal fibroblast showed a different pattern of CGGBP1-CTCF interaction. CGGBP1 and CTCF co-pulldown was positive only when the cells were stimulated after 48h of starvation. The interaction between CTCF and CGGBP1 was stoichiometrically different in these cells as the pulldown of CGGBP1 with CTCF antibody was much weaker than that of CTCF using CGGBP1 antibody. **(C)** CGGBP1-CTCF interaction was confirmed in HEK293T cells by reciprocal co-IPs. **(D)** CGGBP1 and CTCF Co-IP assays were performed in cytoplasmic and nuclear fractions of HEK293T cells separately. CTCF-CGGBP1 pulldowns showed presence of a major protein-protein complex and a minor protein-DNA-protein complex. The purity of nuclear and cytoplasmic fractions were ascertained by using histone H3K9me3 and GAPDH respectively as markers (Figure S2A).

Using published ChIP seq datasets for CGGBP1 (Agarwal et al., 2016) and TFs (Davis et al., 2018) we measured the proximity between binding sites of CGGBP1 and TFs (all identified in repeat-masked hg38). The topmost three transcription factors, binding sites of which most frequently occured in proximity of CGGBP1 binding sites included CTCF and CEBPB, a protein that binds to sites having proximity to CTCF binding sites (Figure S1A and S1B) (Chatterjee et al., 2014; Chin-Tong Ong, 2008; Lefevre et al., 2008; Schmidt et al., 2010). We have earlier shown that CGGBP1 binding sites in Alu repeats match with the binding sites of CEBPB-homolog CEBPZ (Agarwal et al., 2016). The proximity between CTCF and CGGBP1 binding sites was further supported by the findings that centrally enriched CTCF motifs were present in CGGBP1 binding sites. Using CTCFBSDB (Ziebarth et al., 2012) 56% of CGGBP1 peaks showed a centrally enriched presence of a CTCF motif with score >3.0. In addition, L1 and Alu derived 8 bp sub-sequences were also found in CTCF as well as CGGBP1 peaks obtained from repeat-masked alignments (Figure S1, C and D and table S3). CTCF-CGGBP1 occupancy at neighboring sequences would allow functional dependency between the two proteins. Based on these findings we hypothesized that genomic occupancy of CTCF could be affected by the levels of CGGBP1.

### CTCF occupancy preference for repeats over motifs depends on CGGBP1

To understand how CGGBP1 affects genome-wide occupancy of CTCF, we performed CTCF ChIP-seq in HEK293T cells with different levels of CGGBP1. HEK293T cells were transduced with following shRNA lentiviruses: non-targeting (CT), CGGBP1-silencing (KD), CGGBP1-overexpressing (OE) (Figure S2B). We mapped the quality-filtered sequencing reads to repeat-masked hg38 (Table S1) and called peaks. The CT, KD and OE peaks were consistent with previously described CTCF binding sites in HEK293 (ENCFF183AAP, not shown) and other cell types (Figure 3A) suggesting that the overall CTCF occupancy at robust CTCF binding sites are not affected by CGGBP1 levels (Figure 3A).

**Figure 3:**
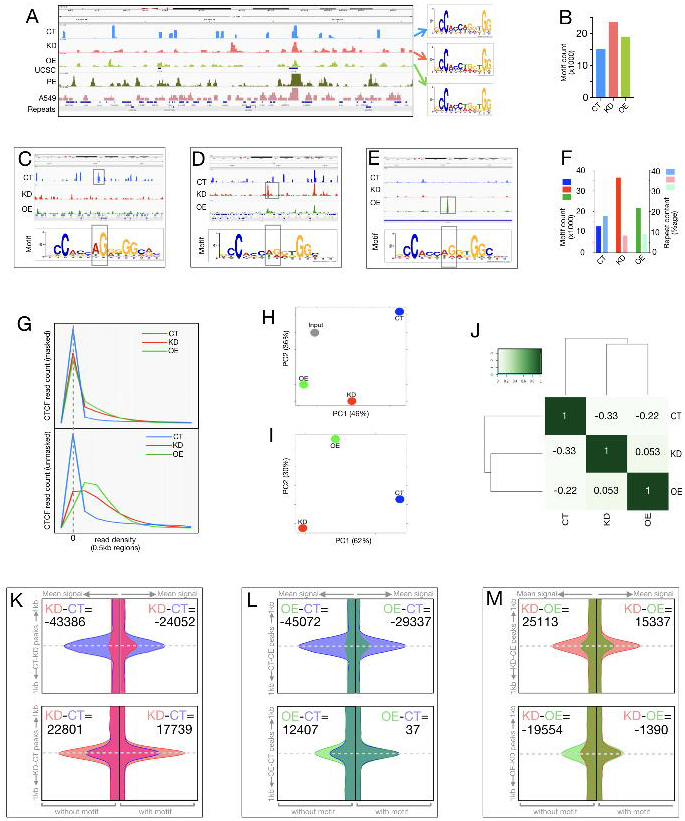
CTCF occupancy at CTCF-binding site motifs and repeats depends on CGGBP1. **(A)** Genome browser tracks showing repeat-masked (RM) CTCF-reads distribution in CT, KD and OE samples in a region on chromosome 21. For comparison, UCSC CTCF-binding sites, CTCF reads from prostate epithelial cells (ENCFF098DGZ) and A549 (ENCFF9810JS) are shown. Widespread differences between cell types and similarities between CT, KD and OE can be seen. The motifs discovered in CT, KD and OE are shown on the right. **(B)** CTCF occupancy at the indicated motifs (A) is low under normal levels of CGGBP1 with KD causing a string increase. **(C to E)** Genome browser tracks showing CTCF read distribution at repeat-unmasked CTCF peaks exclusive for CT (C), KD (D) and OE (E). CTCF peaks and subtle difference in CTCF-motifs enriched in each dataset is highlight in grey box. **(F)** CTCF motif counts and repeat content in repeat-unmasked CTCF peaks were plotted for CT, KD and OE samples respectively. Between CT and KD, a striking shift in CTCF occupancy from repeats to motifs is seen. **(G)** Distribution of repeat-masked (top) and repeat-unmasked (bottom) CTCF reads in randomly picked 0.5kb long 1 million genomic regions for CT, KD and OE samples. CT shows a bipolar distribution pattern with preponderance of read-free and highly read-rich region with a paucity of regions with moderate read density strongly when repeats are unmasked (bottom) as compared to repeat-masked (top). On the contrary, including the repeats shifts the KD and OE read distribution patterns towards the centre with a majority of moderate read density regions. **(H)** Principal Component Analysis (PCA) to find the patterns of differences between CT, KD, OE and input (upper panel) shows that all the three ChIP samples differ from input in different ways. The specificity of CTCF ChIP causes a difference from input that is majorly PC1 for CT, majorly PC2 for OE and a mix of PC1 and PC2 for KD. **(I)** PCA analysis between the ChIP samples only to find the patterns of differences in CT, KD and OE datasets due to CGGBP1 levels showed that CT was nearly equidistant and different from KD (majorly PC1) and OE (majorly PC2) with no similarity between KD and OE. **(J)** Clustering and correlation heatmap for CT, KD and OE samples show that CT is weakly inversely correlated with KD and OE with no correlation between KD and OE. **(K to M)** CTCF reads signal was plotted for CT (blue), KD (red) and OE (green) in 1 kb flanks of peaks centre. Differential peaks as described below were split into without motifs (left flanks) or with motifs (right flanks) and plotted separately. The peak identities are mentioned on the left of each block. The sample from which signal is derived is mentioned in the inset for area under the curve difference calculation. Peak identities are as follows: CT positive KD negative peaks (CT-KD), KD positive CT negative peaks (KD-CT), CT positive OE negative peaks (CT-OE), OE positive CT negative peaks (OE-CT), OE positive KD negative peaks (OE-KD), KD positive OE negative peaks (KD-OE). AUC differences show that repeat-containing motif-free peaks are specifically enriched in CT the most, and the differences between CT, KD and OE are lowest at motif-containing peaks.

A total of 26635, 24418 and 21071 peaks were called in CT, KD and OE respectively (Table S2). The peaks were rich in the CTCF motifs (Figure 3B). MEME on CT, KD and OE peak sequences returned only one motif that corresponded to the HoCoMoCo CTCF-motif (e value limit 3.8e-147). Interestingly, the number of CTCF motifs per peak was lowest in CT, which increased upon knockdown and overexpression of CGGBP1 (Table S2). These findings established the validity of our CTCF ChIP-sequencing assays.

Since these peaks were called using reads aligned only to non-repeat parts of the genome, any occurrence of repetitive sequences was unexpected. However, supporting the observations made on repeat-masked CGGBP1 and CTCF peaks, we observed that even the repeat-masked CT, KD and OE peaks contained LINE-1 and Alu-derived sub-sequences (Figure S1C, S1D and table S3). The occurrence of these motif-like L1 and Alu-derived sub-sequences was strongly enhanced in KD. Hence we mapped CTCF ChIP-seq reads to repeat-unmasked genome and asked if CGGBP1 levels affect CTCF occupancy at repeats.

We next mapped the CT, KD and OE reads to unmasked hg38 and analyzed the CTCF binding patterns (Figure 3, C to E). Even after including the repeats in mapping and peak calling, the only motifs discovered were the canonical CTCF-binding motifs (Figure 3, C to E). Interestingly, the central purine-rich segment (the 7^th^ and 8^th^) of the motifs was different in CT than in KD and OE (regions in the motifs highlighted in figure 3, C to E). A comparison of mappability differences of CTCF bound sequences in repeat-masked and unmasked hg38 is provided in table S4. Interestingly, the peaks that we identified by mapping to unmasked hg38 also showed a very specific association with sequence reads of ENCODE CTCF ChIP seq datasets ENCFF098DGZ and ENCFF9810JS (Figure S3). This meant that the peaks identified on unmasked hg38 are genuine binding sites for CTCF that include repeats. These unmasked CT peaks contained maximum interspersed repeats and least number of CTCF motifs compared to those of KD and OE (table S5 and S6). CGGBP1 knockdown and overexpression reduced the repeat-binding and enhanced motif-binding of CTCF respectively (Figure 3F). The reads appeared relatively dispersed in KD and OE and clustered in CT. A comparison of the dispersion of reads in units of 0.5kb showed that the number of read-free regions are higher in CT than in KD or OE. In KD and OE both, the frequency of zero read density is reduced and the distribution curves shift centrally towards moderate read density (Figure 3G). Interpreted with data shown in figure 3G, it meant that the weak or transient binding sites for CTCF (at which no binding is captured in CT), stabilize in KD and OE thereby reducing the frequency of zero read density regions and increasing the frequency of low read density regions.

CGGBP1 depletion and overexpression both caused a redistribution of CTCF binding. To further understand how do the samples CT, KD and OE differ from each other, we measured differences between the samples using principal component analysis. We analyzed the variation between the CTCF peaks by using input as a control. We could establish that the two principal components accounted for a total of 82% of variation between the samples. The major component (PC1) separated KD and OE in two different direction from the input (located at zero distance from itself). The second component (PC2) clustered the KD and OE closer to each other recapitulating the read density similarity between them (Figure 3H). A second PCA between the ChIP samples (without the input) showed that CT, KD and OE were equidistant from each other such that KD and OE differed from CT in two different directions implying that even if the read densities are similar between OE and KD, the patterns of CTCF binding differs between them (Figure 3I). Hierarchical clustering put the uncorrelated samples KD and OE in one unrelated cluster separate from the inversely correlated sample CT (Figure 3J). Thus, overexpression and knockdown of CGGBP1 showed two different and opposing effects on CTCF binding.

To further understand the nature of differences in CTCF occupancy between CT, KD and OE, we compared read distributions in 1kb flanks of peaks exclusive to each sample using one sample as a negative control at a time. In CT positive KD negative peaks, the background occurrence of KD reads was higher in motif-containing peaks (delta AUC KD-CT= −24052 for motifs and −43386 for repeats) (Figure 3K), whereas the specific presence in CT and absence in KD was tightly associated with repeat-containing reads. In KD positive CT negative peaks the background CT reads were present in peaks with and without motifs both (a much reduced delta AUC KD-CT = 17739 for motifs and 22801 for repeats) (Figure 3K). These findings meant that CGGBP1 mitigates the binding of CTCF at repeat-free motif-rich peaks and knockdown of CGGBP1 led to an enhancement of this weak binding in motif-rich KD positive CT negative peaks. Interestingly, compared to the CT positive OE negative peaks, the OE positive CT negative peaks showed a different kind of change in read distribution. OE positive CT negative peaks had an increase in OE-specific reads only at peaks without motifs with negligible difference at motif-containing peaks (Figure 3L). A comparison of KD-CT and OE-CT delta AUC values show that KD is a stronger driver of CTCF occupancy at CTCF motifs whereas OE drives CTCF occupancy at regions which are repeat-rich and motif-poor. Finally, a reciprocal comparison between OE and KD exclusive peaks confirmed this again (Figure 3M). While KD exclusive peaks showed higher CTCF binding at regions with and without motifs both, the OE exclusive peaks showed enhanced CTCF binding at regions without motifs only (a much reduced delta AUC KD-OE = −1390 for motifs and 19554 for repeats). An interpretation of these results in the light of repeat contents of the motif containing and motif free exclusive peaks (Table S7 and S8) establishes that (i) CTCF binds to high-repeat low-motif as well as high-motif low-repeat regions, (ii) knockdown of CGGBP1 shifts the binding from repeats to the motifs, and (iii) overexpression of CGGBP1 exerts an opposite effect and shifts the binding to repeat-rich regions. These results also showed that CGGBP1 levels determine CTCF binding pattern at repetitive sequences.

Consistent with the above findings, measuring of repeat content in CT, KD and OE peaks showed that CTCF binding sites were excessively motif rich in KD and OE and motif-poor but L1-rich in CT (Table S5). These findings showed that although repeat-masked CT, KD and OE peaks all have CTCF motif as the consensus binding site, after repeat masking this motif is retained strongly in KD and eliminated from the CT peaks. Thus it seems that CTCF normally binds to repeats and motifs both but when CGGBP1 levels are disrupted, the binding is adversely affected at repeats. This shifts the balance of of CTCF occupancy towards motif-richness in KD.

Identification of CGGBP1 binding sites at repeat-masked regions also showed that CGGBP1 binding sites are associated with multiple small L1-derived motifs that could not be identified as repeats by RepeatMasker (Figure S1C and S1D). We argued that if CGGBP1 and CTCF binding sites occur in proximity with each other (as shown in Figure S1) then CGGBP1 binding sites will occur with same frequency in the neighbourhood of CTCF binding sites that are L1-rich in CT and motif-rich in KD and OE. Indeed the short L1-derived motifs were found with comparable frequencies in CT, KD and OE peaks despite the differences in their L1 and motif contents (Table S9).

With these findings we concluded that CGGBP1 regulates the binding pattern of CTCF through the proximity of L1-derived CGGBP1 binding sites to CTCF-binding sites. Next we wanted to identify the functional effects of the changes in CTCF binding pattern upon changes in CGGBP1 levels.

### CGGBP1 level affects CTCF occupancy at known insulators

For finding out the effects of altered CTCF binding caused by changes in CGGBP1, we first analyzed the disturbances in CTCF occupancy at genomic locations annotated as regulatory elements (UCSC Regulation datasets). CTCF binding at the regulatory elements influences genome organization and function with direct effects on epigenetic state of the chromatin, transcription, replication and genomic integrity. Read density measurements at regulatory regions showed strong enhancement of CTCF binding at replication origins (Figure S4A) and enhancers (Figure S4B) by CGGBP1 overexpression.

Topologically associating domains have been described for HEK293 cells (Zuin et al., 2014). We analyzed the CTCF occupancy at these topologically associating domains (TADs) in CT, KD and OE. CGGBP1 depletion caused a loss of CTCF binding at 1128 TADs and a gain of CTCF binding at 390 TADs. These findings suggested that new TADs are formed in KD due to CTCF binding site rearrangements. The Lamina Associated Domains (LADs), identified in Tig3ET cells (Guelen et al., 2008), are conserved between different cell types. The LADs showed a consistent reduction in CTCF binding at LAD start and end sites in KD as compared to CT and OE (Figure 4A and 4B). The LADs are rich in L1 repeats (van Steensel and Belmont, 2017) and thus a loss of binding in KD was expected.

**Figure 4:**
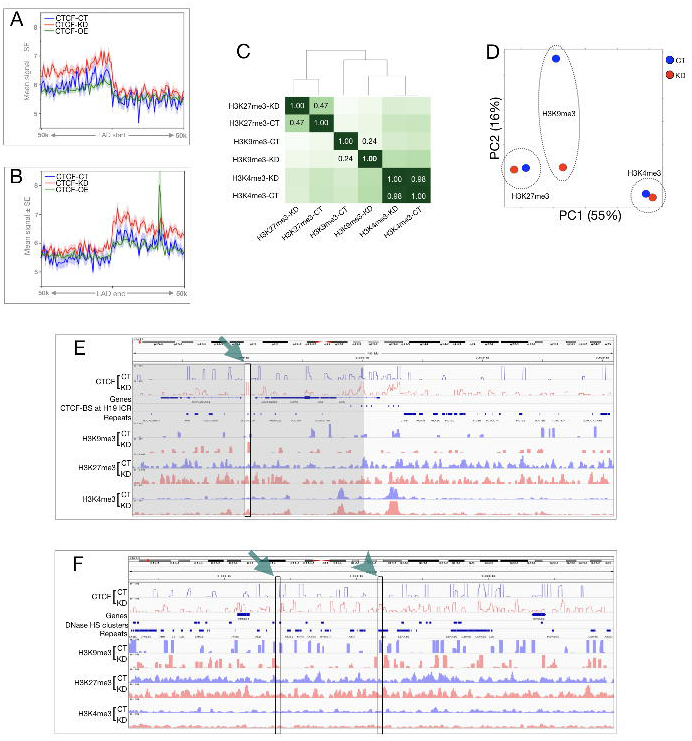
CGGBP1 affects CTCF occupancy and histone modifications at known CTCF-binding sites. **(A** and **B)** CTCF read distribution was plotted for CT, KD and OE at the LAD boundaries in the 50kb flanks with bins of 1kb. The average read density with standard error of mean were plotted for LAD-start sites **(A)** and LAD end sites **(B)**. CTCF binding is reported to occur in 10 kb flanks of the LAD start and end sites. **(C)** Correlation-heatmap was plotted for H3K4me3, H3K9me3 and H3K27me3 ChIP-seq peaks in CT and KD. The correlation values (r) corresponding to CT and KD are mentioned for each histone modification in the heatmap. Unlike H3K4me3 and H3K27me3, the CT and KD samples for H3K9me3 were significantly different from each other. **(D)** PCA analysis using input as control shows the distance between H3K9me3 CT and KD as the most affected in KD. CGGBP1 depletion minimizes the distinction of H3K9me3 from H3K4me3 and H3K27me3 along PC2 only while along PC1, the two repressing marks remain closer to each other than the activating mark. (**E)** Disturbances in the H3K9me3 is observed in KD compared to CT. Genome browser tracks showing reads of CTCF, H3K9me3, H3K27me3 and H3K4me3 for CT and KD along with the CTCF-binding sites and repeats at the H19 ICR locus. The gain of CTCF-binding upon CGGBP1 knockdown is highlighted by green arrow and changes in the histone modification profile in 10 kb flanks are highlighted by grey boxes. **(F)** Genome browser view of the beta globin locus control region undergoing change in chromatin barrier function upon CGGBP1 depletion. The plot contains tracks of CTCF reads, H3K9me3, H3K27me3 and H3K4me3 reads, hg38 genes, DNase hypersensitivity clusters and repeats. The gain and loss of CTCF binding upon CGGBP1 loss-of-function are highlighted with an arrowhead and an arrow respectively.

Although the OE system is of value for testing the dependence of CTCF DNA binding on CGGBP1 levels, we argued that it is more artifactual than KD for studying the functional outcome of disruption in CGGBP1-dependent CTCF-DNA binding. Hence, we analyzed the effects of KD on CTCF occupancy at regulatory elements by comparing it with those of CT. These analyses were complemented with genome-wide measurements of three different histone modifications in CT and KD samples: H3K4me3, H3K9me3 and H3K27me3 (Table S10). Unsupervised clustering and PCA of the three histone modification patterns in CT and KD showed that CGGBP1 knockdown disrupted H3K9me3 the most, with only minor effects on H3K4me3 and H3K27me3 (Figure 4C and 4D).

Of all the regulatory regions that we analyzed, we observed changes in CTCF occupancy accompanied by histone modification changes at LADs and CTCF-binding sites (motif-dependent sites located in repeat free regions, called UCSC CTCF sites). CTCF loss of occupancy at LADs was accompanied by a mild but consistent loss of the repeat-silencing mark H3K9me3 (Figure S5A and S5B). At UCSC CTCF sites, the gain of CTCF binding in KD was accompanied by strong increases in transcription activating histone mark H3K4me3 with only mild changes in H3K9me3 and H3K27me3 marks (Figure S6A). However, the UCSC CTCF sites which are repeat-poor and CTCF motif-rich had maximum differences between CT and KD for H3K4me3 occupancy (Figure S6A and S6B). H3K9me3 is a histone mark that identifies repetitive sequences and we further investigated if differences in CTCF occupancy and H3K9me3 between CT and KD occur predominantly at repeats and repeat-derived regulatory regions.

The insulators in the beta globin locus and H19-ICR are known CTCF binding sites. We analyzed the patterns of CTCF binding in CT and KD at these two candidate regions. At the three CTCF binding sites located upstream of H19, the most distal site upstream of the TSS showed the strongest binding of CTCF was observed on. The three upstream binding sites, including the intervening regions between them, showed a conspicuous increase in CTCF binding in KD. This change in the pattern of CTCF-binding was also concomitant with a change in chromatin marks from bivalent (co-occurrence of H3K9me3 and H3K4me3) in CT to active (only H3K4me3) in KD due to a loss of H3K9me3 (Figure 4E). Similar changes were observed at the downstream CTCF-binding sites where CGGBP1 knockdown led to an increase in CTCF-binding. Further downstream from the H19 gene, there were distinct sites of gain and loss of CTCF-binding in KD. A comparison of histone marks in 10 kb flanks showed that the asymmetry of H3K9me3 maintained on two sides of a binding site in CT was lost upon disruption of CTCF binding in KD (arrow). One such region with a gain of CTCF binding in KD causing loss of H3K9me3 differences in the flanks is highlighted in figure 4E under shadows.

In the beta globin LCR, CTCF knockdown caused a disruption of binding such that gain (arrowhead) as well as loss (arrow) of binding in KD was observed at various DNAse hypersensitivity sites. The H3K9me3 signal in 10kb flanks of these sites was symmetric in CT and became asymmetric in KD. Although there were changes in H3K27me3 and H3K4me3 signals as well, but a specific change in the region lying between loss and gain of CTCF binding sites in KD was observed only for H3K9me3 (Figure 4F).

Of the two candidate regions analyzed here, H19 ICR is relatively poor in repeats than the beta globin LCR and H19 ICR region also showed a weaker disturbance in H3K9me3 marks than beta globin LCR region. These findings suggested that the change in the patterns of CTCF caused by CGGBP1 knockdown occupancy can affect chromatin barrier functions. It is also indicated that this CGGBP1-regulated chromatin barrier function through CTCF is predominant at repeats and thus affects the repeat-marking histone modification H3K9me3.

### CGGBP1 regulates chromatin barrier activity at motif-independent CTCF-binding sites in repeats

Next, we analyzed the gain or loss of chromatin barrier functions at regions that undergo loss or gain of CTCF binding upon CGGBP1 knockdown. We hypothesized that repeat-rich regions that have CGGBP1-dependent CTCF binding sites function as CGGBP1-dependent chromatin barrier sites as well. To test this possibility, we first identified CTCF binding sites in CT and KD that we could unambiguously classify as motif-free repeats or repeat-free motifs (Figure 5A). Any CGGBP1-dependent change in CTCF binding and hence on chromatin barrier function would be limited to repeat containing CTCF binding sites that differ between CT and KD. By comparing CT-KD common peaks against CT/KD exclusive peaks, we sought to identify if indeed CGGBP1 selectively disrupts barrier function of those CTCF binding sites that are motif-free repeats (Figure 5B).

**Figure 5:**
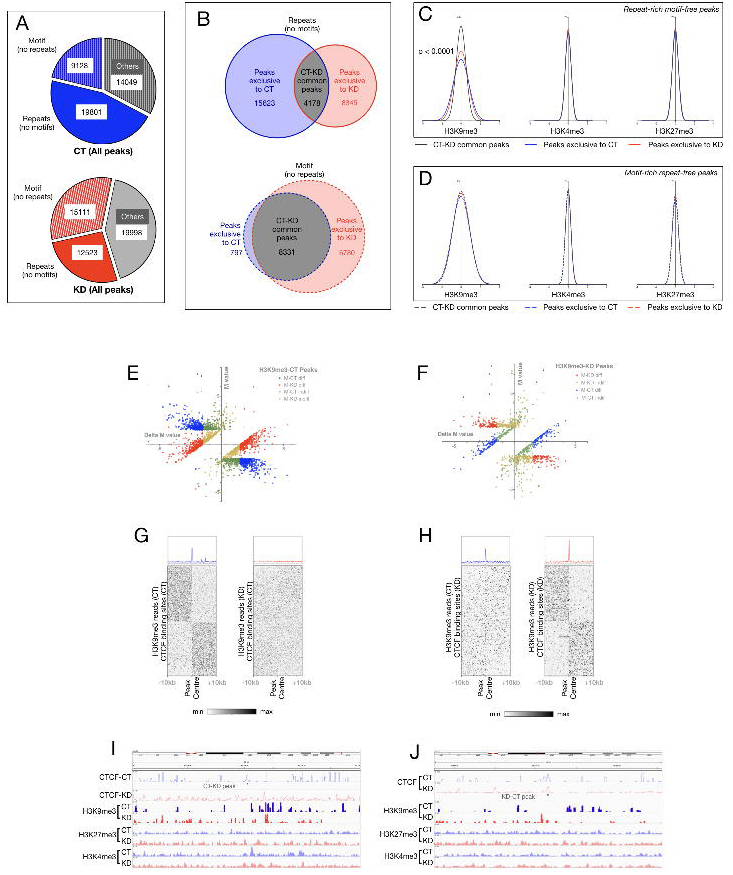
Changes in H3K9me3 occupancy upon CGGBP1 knockdown are tightly linked to motif-free repeat-rich CTCF-binding sites. (A) All CT (blue, top) and KD (red, bottom) peaks were classified into motif (no repeats) or repeats (no motifs) using FIMO and RepeatMasker tools. Peaks that could not be unambiguously classified were labelled as Others (grey) in both the groups. (B) Peaks exclusive to CT (red) or KD (blue) were determined in subsets of CT and KD peaks shown in A. Venn diagrams depicting shared or exclusive peaks between 9128 striped blue and 15111 striped red (shown in A) are shown on top in B. Venn diagrams depicting shared or exclusive peaks between 19801 solid blue and 12523 solid red (shown in A) are shown on bottom in B. These subsets of peaks help comparing CT and KD for features that vary between them depending on repeat or motif contents exclusively. (C and D) Frequency distribution of M values of differences in H3K9me3, H3K27me3 and H3K4me3 read counts (upstream-downstream 10kb flanks for the six sets of peaks depicted in B) in repeat-rich (no motif) peaks(C) and motif-rich (no repeat) peaks (D). Only H3K9me3 shows weak differences of CT and KD exclusives from common peaks for motif-rich (no repeat) peaks (D). H3K9me3 distribution differs strongly between CT and KD and deviates from common peaks at repeat-rich (no motif) peaks (C) only. (E and F) The M values depicted in H3K9me3 datasets (D) were separately analyzed for CT and KD to filter out peaks 10 kb flanks of which satisfy two conditions: (i) M value upstream-downstream difference >1.5 and (ii) the deltaM [(M of KD) - (M of CT)) >1.5 (E; red and blue spots) or deltaM [(M of CT) - (M of KD)) >1.5 (F; red and blue spots). (G and H) Heat maps of H3K9me3 signals from CT and KD datasets in the peaks corresponding to red-blue peaks in E and H show that there is a clear difference in H3K9me3 signal transitioning exactly at the peak centre where CTCF binding is observed and affected by CGGBP1 in KD. F and G show peaks exhibiting loss and gain of barrier activity respectively. (I an J) Genome browser tracks showing loss of CTCF-binding at a representative CT-KD peak that also exhibit loss of H3K9me3 barrier function upon CGGBP1 knockdown (I). A representative KD-CT exclusive peak that show gain of CTCF binding upon CGGBP1 knockdown also displayed gain of H3K9me3 barrier function (J).

For CT-KD common peaks and CT/KD exclusive peaks, the differences in H3K4me3, H3K9me3 and H3K27me3 read counts in 10 kb upstream and downstream flanks were compared. Motif-free repeat-containing CT-KD common peaks were sites with a lesser barrier function as a distribution of the difference between upstream and downstream H3K9me3 signals at such peaks showed a low deviation from the normal of zero difference. The motif-free repeat-containing CT/KD exclusive peaks on the other hand showed a broader distribution with significantly higher deviation from the normal (Figure 5C). The distribution of H3K9me3 upstream-downstream difference for the CT exclusive motif-free repeat-containing peaks were the most different from those of the CT-KD common peaks and were also significantly different from those of the KD exclusive motif-free repeat-containing peaks. Strikingly, in the motif-containing repeat-free category, the H3K9me3 upstream-downstream difference distribution patterns of CT/KD exclusive peaks were not different from the common ones (Figure 5D). Furthermore, H3K4me3 and H3K27me3 upstream-downstream difference showed no difference between CT/KD exclusive peaks and CT-KD common peaks (Figure 5C and 5D). These findings demonstrated that H3K9me3 occupancy in the flanks of motif-free CTCF-binding sites in repeats are regulated by CGGBP1. Gain as well as loss of barrier function is observed at such regions upon CGGBP1 knockdown as CT-KD as well as KD-CT peaks both showed significantly different upstream-downstream H3K9me3 signals compared to the CT/KD common peaks (Figure 5C and 5D). Changes in barrier activity due to altered CTCF binding is the most likely reason for disruption in the H3K9me3 patterns. These regions could be potential insulator sequences regulated by a CGGBP1-CTCF crosstalk.

To identify such regions, we selectively fished out the regions with strong gain or loss of CTCF binding (CT/KD exclusive peaks) that also exhibit a strong change in H3K9me3 occupancy in 10kb flanks. We applied arbitrary threshold (Figure S7) to select a subset of exclusive peaks that showed strong differences in upstream and downstream signals (Figure S7). Thus, upon CGGBP1 knockdown, the largest change in histone modification landscape occurs for H3K9me3 in the flanks of sites with CGGBP1-dependent CTCF binding. These potential chromatin boundary sequences were of two kinds: those which lost the normal binding in KD (Figure 5E-G) and those with gained *de novo* binding in KD (Figure 5H-J). In comparison, CT/KD exclusive peaks with above threshold changes in H3K4me3 and H3K27me3 occupancy in 10kb flanks were much fewer in number as well as with lower upstream-downstream differences (Figure S8). Thus, we identified 663 sites with a loss of barrier function and 216 sites with a gain of barrier function upon CGGBP1 depletion and a concomitant loss of CTCF occupancy.

These potential chromatin boundary elements with a loss of binding in KD (Figure 5E and 5G) were most repeat-rich (35% L1 content), whereas those with a gain of binding in KD (Figure 5F and 5H) were repeat poor and highly CTCF motif rich (Table S11). Thus, we concluded that CTCF binding sites located in repeats or in repeat-proximity function as barriers between opposing patterns of H3K9me3. CTCF binding to such sites and the barrier function of these sites both depend on CGGBP1. At the same time, there are sites to which de novo binding of CTCF takes place in the absence of CGGBP1. These anomalous CTCF-binding sites exhibit barrier activity in the absence of CGGBP1. Changes in CGGBP1 levels seem to alter the chromatin barrier function of repeat-derived candidate insulator sequences through regulation of CTCF binding.

## DISCUSSION

Human CTCF is a multifunctional protein pivotal to the functional organization of chromatin (Ong and Corces, 2014). Some well established functions of CTCF are regulation of insulators and boundary elements, regulation of topologically associating domains and higher order chromatin structure (Ong and Corces, 2014). CTCF in complex with proteins including cohesin ring members orchestrates chromatin structure and changes in CTCF binding has been shown to affect histone modifications and depend on cytosine methylation (Maurano et al., 2015; Ong and Corces, 2014; Wang et al., 2012). Given the important functions executed by CTCF, it makes evolutionary sense that there are regulatory crosstalks between CTCF and other proteins. Of the nearly 70 protein interactants of CTCF (NCBI Gene), the mechanisms of interactions and functional crosstalks between only some have been worked out. Some notable examples are SMCs (Stedman W), YY1 (Weintraub et al., 2017), BRD proteins (Hsu SC; Weintraub et al., 2017) and NPM (Yusufzai et al., 2004). Identification of new interactants and regulators of CTCF will lead to a deeper understanding of chromatin regulation and function. The work presented here shows that human CGGBP1 is a regulator of CTCF occupancy and through it, it also regulates chromatin barrier activity of repeats.

The colocalization of CTCF with CGGBP1 in the nucleus is unsurprising as both proteins have high concentration in the nucleus. However, colocalization at the only extranuclear site of CGGBP1 and CTCF expression, midbody, suggests that functional crosstalk between these two proteins might play a role in cytokinesis, abscission checkpoint and regulation of ploidy. Repetitive sequences prone to faulty homologous recombination can lead to chromosomal fusions (Boissinot et al., 2006; Robberecht et al., 2013; Sasaki et al., 2010). Interestingly, the function of repeat-binding proteins such as CGGBP1, which delay cytokinetic abscission to resolve internuclear chromatin bridges, may cooperate with other midbody proteins for genomic integrity. The fact that CTCF is present at midbodies leads to an interesting possibility that similar to CGGBP1, CTCF may also contribute to genomic integrity through mechanisms similar to those of CGGBP1 (Kemp et al., 2014). In line with this possibility, we have observed regulation of binding of CTCF at repeats by CGGBP1.

We have shown the interactions between CTCF and CGGBP1 through co-immunoprecipitation and PLA on endogenously expressed proteins at native levels. This assumes significance because co-immunoprecipitation studies using overexpression systems are prone to detecting interactions that may not hold true at the much lower physiological concentrations of the two proteins (Moriya, 2015). We have rigorously tested the interactions in multiple cell lines and have established that using targets other than CTCF which are also expressed at high levels in nuclei, we fail to detect an interaction in PLA (not shown). PLA results, which showed that most of the CTCF-CGGBP1 interactions occur in the nuclei, set the ground for the working hypothesis of the present work that the interaction between CGGBP1 and CTCF is required for proper DNA-binding and DNA-binding-dependent functions of the two proteins. There are supporting evidences for this possibility in the literature. The most direct evidence is the YY1-CTCF complex study in which CGGBP1 emerged as a co-player with CTCF and YY1 for insulator and enhancer activity regulation (Weintraub et al., 2017). The DNA binding of CGGBP1 in the vicinity of CTCF and YY1 leads to a possibility that in addition to a direct protein-protein complex, some interaction between these proteins could be mediated through a DNA bridge. Indeed our co-immunoprecipitation assays in nuclear fractions with DNAse digestion prove that a fraction of the CTCF-CGGBP1 co-immunoprecipitates is bridged by DNA. Since the c-immunoprecipitations were performed in cleared lysates with long DNA molecules pelleted out and discarded, it implies that the DNA-bridged complexes of CTCF and CGGBP1 are not artefactual interactions mediated by long DNA molecules. However, the DNA-bridged interactions only account for approximately 34% of the total complex. Given that CGGBP1 binds to repetitive DNA, one interpretation of our co-immunoprecipitation findings is that the larger protein CTCF binds to a set of specific sites on the repetitive DNA whereas the small protein CGGBP1 binds to the same sequences through a physical complex with CTCF as also in the immediate vicinity of the CTCF-bound DNA. Thus CGGBP1 could restrict CTCF binding to specific subsequences within the repeats by acting as a border marker for CTCF binding sites at repeats. Our ChIP-seq results favour this possibility.

We chose to study CTCF-CGGBP1 crosstalk in HEK293T cells because these cells have expression of both the proteins strong enough to reason that a knockdown of CGGBP1 will have significant effects. In addition to having strong native expression of CGGBP1, HEK293T cells showed an ability to tolerate a large change in levels of CGGBP1, from a near-complete knockdown to approximately 7.70 folds overexpression. This we have not observed in any other cell type in which CGGBP1 has been studied before. Unfortunately, CTCF ChIP-seq data from HEK293T are not available (except for the dataset ENCFF183AAP on ENCODE that is flagged for low read count) to compare our results with. The CTCF ChIP-seq in HEK293T we have performed shows a robust correlation with the CTCF ChIP-seq data available from other cell lines thereby establishing the specificity of CTCF pulldown in our assays. This comparison with validated CTCF ChIP-seq datasets is necessary because we have mapped CTCF-bound DNA sequence reads to human genome with as well as without repeat masking. By retaining only the best and uniquely mappable reads, we have ensured that any cross-matching of reads to repeat sequences were minimized to the best possibility. Both the repeat-masked and repeat-unmasked aligned read sets gave rise to distinct sets of peaks. The repeat-masked peaks were rich in the canonical CTCF motif (Kulakovskiy et al., 2018), whereas the repeat-unmasked peaks were repeat-rich and relatively motif poor. The CTCF-binding peaks from the published CTCF ChIP-seq datasets showed concentration of reads from our CTCF peaks obtained from repeat-unmasked (L1 rich and motif poor) as well as repeat-masked alignments (motif rich and L1-poor). This indicated that the CTCF peaks called in previously published studies (Wang et al., 2015) have not included the CTCF-binding sites at repeats and focussed on peaks that contain CTCF-binding motifs. Interestingly, the CTCF-motif we observed in repeat-masked datasets were also seen in repeat-unmasked datasets albeit at a lower frequency. This suggests that the binding of CTCF to repeats does not occur on consensus sequences that correspond to any specific sequence motif. Given that the preference for CTCF occupancy at repeats was dependent on CGGBP1, it seems that there is a cooperative association between CTCF and CGGBP1 for binding to the repeats or on CTCF-motifs occurring within or in immediate proximity to the repeats. This would also justify the findings that CTCF-binding sites have evolved out of repeats, especially retrotransposons (Schmidt et al., 2012). Our findings that Alu and L1-derived subsequences that were prevalent in CGGBP1 binding sites were also seen at high frequency in CTCF-binding sites lends further support to this argument.

One of the most important aspects of our findings is that in presence of normal levels of CGGBP1, the CTCF-binding sites showed very strong pileups of reads with sharp transitions into read-free regions. In comparison, upon CGGBP1 knockdown the binding pattern of CTCF became more centrally enriched in reads with a normally distributed read pattern. The gradient ends of CTCF-binding sites in KD as compared to blunt ends in CT suggest that CGGBP1 restricts and stabilizes CTCF binding to target sequences. This is reflected in the higher number of reads per peak in CT compared to KD, without an increase in peak length. Presence of CTCF-binding sites inside and in the vicinity of repeats makes this more plausible because binding of CGGBP1 to repetitive sequences flanking the CTCF target sites and motifs can constrain the binding of CTCF. A comparison of CTCF-binding sites (Estécio et al., 2012; Jordà et al., 2017; Kim, 2008) and CGGBP1-binding sites in fact shows proximity as well as an overlap at L1 and Alu repeats. In addition, proximity between CGGBP1 and CTCF binding has been shown at the CTCF-bound enhancer-promoter loops (Weintraub et al., 2017). CGGBP1 thus seems to exert a qualitative effect on CTCF-DNA binding. CGGBP1 also regulated CTCF occupancy on LADs, TSSs and the sets of exclusive peaks that we have identified. CGGBP1 forms complex with NPM1 (Yusufzai et al., 2004) and the effects on LADs could be an outcome of CTCF-NPM1-CGGBP1 crosstalk at lamina associating sequences, often rich in L1 retrotransposons (van Steensel and Belmont, 2017). Interestingly, overexpression of CGGBP1 changed CTCF binding pattern in two different ways. At most regions, it appeared similar to KD with a loss of sharp transitions from CTCF-bound to CTCF-unbound regions. At sites where CGGBP1 regulates cytosine methylation patterns such as replication origins and enhancers (Patel et al., 2018), the overexpression of CGGBP1 caused a drastic increase in CTCF-binding. At the same regions, an opposite effect of loss of CTCF binding was observed in the absence of CGGBP1 in a range of 1kb. Thus KD and OE exert opposite effects on CTCF occupancy at loci where CGGBP1 regulates cytosine methylation. However, the similarities between CTCF-binding pattern in two contrasting states of CGGBP1 knockdown and overexpression could be due to many reasons. CGGBP1 is required for CTCF binding to repeats and yet CGGBP1 itself is a repeat binding protein. Excess amounts of CGGBP1 could potentially interfere with CTCF-binding to repeats. CGGBP1 is a growth signal sensor protein that binds to the DNA depending on post-translational modifications (Agarwal et al., 2016). The phosphorylation by ATR at S164 and RTKs on Y20 seem to be essential for its nuclear localization and DNA binding. In the absence of signals that generate optimal post-translational modifications on CGGBP1, we have seen that its overexpression exerts a dominant negative effect (Agarwal et al., 2016; Singh et al., 2014). This is why the overexpression of WT CGGBP1 with insufficient post-translational modifications seems to mimic the effects of its knockdown partially. In addition, CGGBP1 itself is a repeat-binding protein and in OE excess of CGGBP1 can compete for repeat occupancy with CTCF and effectively negate the CGGBP1-dependence of CTCF for repeat binding. Since overexpression systems are prone to artefacts due to toxicity of overexpression, we have restricted the use of the OE sample only to establish that CTCF-DNA binding depends on levels of CGGBP1. To study the functional outcome of CTCF dependence on CGGBP1, we restricted only to the CT and KD samples. These two samples allowed us to study the effects of CGGBP1 on CTCF without altering the stoichiometry of CGGBP1 limitlessly in an overexpression system. Thus in the light of our findings that insulator sites in H19 and beta globin loci underwent a change in CTCF occupancy upon CGGBP1 knockdown, we restricted our analyses to find out similar changes genome-wide in upon CGGBP1 knockdown.

The H3K9me3, H3K4me3 and H3K27me3 represent three different kinds of histone modifications, transcriptional suppression of repetitive DNA, transcriptional activation (poised state of transcription with co-occurrence of H3K9me3, H3K4me3) and lineage specific silencing of genes in the course of differentiation respectively. CGGBP1 is a repeat-binding protein and changes in repeat silencing modification H3K9me3 as an effect of CGGBP1 knockdown is not surprising. Of high importance is the observation that this change occurs very strongly at 879 locations in the genome with CGGBP1-regulated CTCF-binding sites that we have pulled out by applying stringent thresholds of H3K9me3 signal differences in the CTCF-binding site flanks in CT and KD. At these binding sites, CTCF maintains a certain pattern of H3K9me3 in the flanking regions that depends on CGGBP1. Considering these CTCF-binding sites as boundaries between contrasting histone modification patterns, CGGBP1 emerges as a regulator of chromatin boundary element function of CTCF. By comparing CT with KD, we have identified new locations with gain and loss of boundary element functions. This identification of novel boundary elements is robust as it is based on a three step filtration: binding of CTCF on these sites, CGGBP1-dependent changes in CTCF binding, and changes in H3K9me3 but not necessarily in other modifications asymmetrically in the flanks of these sites. The repeat and motif contents in these sets show that upon CGGBP1 depletion the boundary element-like activity of repeat rich regions yield to repeat-poor and motif-rich CTCF-binding sites. CGGBP1 forms complex with histone methyltransferase SUV39H2 (Stelzl et al., 2005) and a loss of regulation of SUV39H2 by CGGBP1 in KD could enable a selective change in H3K9me3 and not other histone marks. Our analyses reveal that there are repeat elements that act as boundary elements through a CGGBP1-CTCF crosstalk. Genome-browsing of ENCODE datasets does show abundant CTCF binding at repeats however the regulation of this has remained unclear so far and our findings provide some explanations. Thus, most CTCF-bound repeats functioning as boundary elements depend on CGGBP1 whereas boundary element activity of repeat-free sites containing CTCF-binding motif remain unaffected. CGGBP1-CTCF axis regulating boundary element functions of repeats affect histone modifications specifically. Of the three histone modifications we assayed, only the repeat silencing specific histone modification H3K9me3 was altered. Interestingly, CGGBP1 forms complex with histone methyltransferase SUV39H2 (Stelzl et al., 2005). CGGBP1 regulation of CTCF binding could be important for repeat heterochromatinization and silencing. A disruption of repeat heterochromatinization could alter gene expression patterns, and also affect genomic integrity.

In summary, we have discovered CGGBP1 as a regulator of CTCF DNA binding pattern with a direct effect on boundary element like functioning of repeat-rich and motif-rich regions. Our results demonstrate that there is a functional outcome of the CTCF binding preference on repeats or motifs that determines the function of genomic sites as boundary elements. This pivotal function of CTCF depends on CGGBP1. CGGBP1 has evolved later than CTCF. Thus, CGGBP1 is not required for DNA binding of CTCF, but only acts as a fine adjuster of CTCF binding pattern and through it chromatin structure and function. The CGGBP1-CTCF crosstalk is thus an essential part of functioning of CTCF.

## AUTHOR CONTRIBUTIONS

MP and DP conducted experiments, analyzed data and wrote the manuscript. US led the project, analyzed data and wrote the manuscript. MP and DP have contributed equally to the work.

## ACKNOWLEDGEMENTS

The authors acknowledge the help of HoMeCell Lab members in the carrying out of the project and critical reading of the manuscript. The authors are thankful to Gujarat Biotechnology Research Centre for sequencing services, ISTF (IIT Gandhinagar) for support with computational resources and publicly available data, resources and tools for their valuable use in this work. This work was supported by funding from DBT, DST and GSBTM to US.

## METHODS

### Cell culture and lentivirus transduction

Human dermal fibroblast (Sigma, passage 15-24), human juvenile foreskin fibroblast (Himedia, passage 5-30) and HEK293T cells were grown in DMEM (AL007A) supplemented with 10% FBS. Control or CGGBP1-shmiR (targeting 4 different regions in CGGBP1 ORF) or CGGBP1-overexpression lentivirus constructs were obtained from Origene. The third generation lenti-packaging plasmids: pRSV-Rev (12253), pMDLg/pRRE (12251) and pMD2.G (12259) were obtained from Addgene. For lentiviral production, the packaging plasmids and lentiviral constructs were mixed in equimolar ratios and used for transfection. Transfection was performed by using FuGene (Promega). 1:10000 diluted 10 mg/ml polybrene (Sigma) was used for transducing HEK293T cells. For stable transduction control and CGGBP1 shmiR traduced cells were selected with 10 ug/ml puromycin (Himedia).

### Immunofluorescence

CGGBP1 and CTCF immunofluorescence were carried out in human juvenile foreskin fibroblast using standard protocol. In brief, cells were fixed for 10 minutes in 3.7% formaldehyde solution followed by permeabilization with 1% Triton X-100 in PBS for 10 minutes. After permeabilization, cells were incubated with 10% FBS, 0.05% Triton X-100 in PBS for an hour. Subsequently, cells were incubated with primary antibodies (Anti-CGGBP1 rabbit polyclonal and anti-CTCF mouse monoclonal) for 2 hours at room temperature followed by incubation with secondary antibodies (anti-mouse Alexa fluor 594 and anti-rabbit Alexa fluor 488) for 2 hours at room temperature. Samples were counterstained with Fluoroshield Mounting Medium (Ab104135) containing 4′,6-diamidino-2-phenylindole. Samples were examined using a Leica confocal microscope (magnification x630).

### Proximity Ligation Assays (PLA)

Duolink PLA kit was used to detect protein-protein interactions (Sigma Aldrich DUO92101). PLA was performed according to manufacturer’s protocol plus additional negative controls. In brief, Human Juvenile fibroblast were fixed with 4% formaldehyde solution at 37°C for 10 minutes followed by permeabilization with 1% Triton X-100 in PBS for 10 minutes at room temperature. Cells were washed with PBS three times followed by incubation in blocking buffer at 37°C for 1 hour. Samples were subsequently incubated with primary antibody pairs (two pairs of specific antibodies, one pair each of a specific and a non-specific antibody) or only blocking buffer at room temperature for 2 hours. Samples were incubated with oligo probe-conjugated secondary antibodies and ligation and amplification were carried out to produce rolling circle PCR product. Amplified products were detected by hybridization with Texas Red-labeled oligonucleotides and sample were counterstained with Fluoroshield Mounting Medium (Ab104135) containing 4′,6-diamidino-2-phenylindole. Samples were examined using a Leica confocal microscope (magnification x400).

### Co-Immunoprecipitation and immunoblotting assay

Cell extracts were subjected to pre-clearance by incubating with non-specific antibody/IgG and Protein G sepharose beads (GE 17-0618-01). Pre-cleared cell extract was incubated with CGGBP1 and CTCF antibodies separately overnight followed by incubation with Protein G sepharose beads for 2 hours. Protein bound sepharose beads were washed 4 times with PBS. Beads-bound fraction was subjected to elution by boiling in SDS-Laemmli buffer followed by SDS PAGE and immunoblotting with indicated antibodies.

### Co-Immunoprecipitation followed by DNaseI digestion assay

Cytoplasmic and nuclear fraction were separated from HEK293T cells by REAP protocol (Suzuki et al., 2010). Cytoplasmic fraction was pre-cleared by incubating with Protein G sepharose beads followed by immunoprecipitation with CGGBP1 and CTCF separately. Nuclear extracts was pre-cleared by incubating with Protein G sepharose beads. Pre-cleared nuclear extract was incubated with primary antibodies overnight followed by incubation with Protein G sepharose beads for 2 hours. For DNaseI digestion, the immunoprecipitated protein-bound sepharose beads was washed four times with PBS and suspended in the DNaseI digestion buffer containing DNaseI (6 Units of M0303S NEB). The samples were incubated for 10 minutes at room temperature and centrifuged at low speed to separate two fractions: beads-antibody-bound proteins (pellet) and proteins released by DNaseI digestion (supernatant). The separated fractions were subjected to SDS PAGE followed by western blotting (Suzuki et al. 2010).

### CTCF ChIP-sequencing

CTCF ChIP was performed by using MAGnifyTM Chromatin Immunoprecipitation System kit (Invitrogen 49-2024) with minor modifications to the protocol. HEK293T cells were transduced with non-targetting shRNA, CGGBP1-targetting shRNA or CGGBP1-overexpressing lentiviruses. The shRNA-transduced cells were selected with puromycin (10 µg/ml) for 7 days. Approximately 200 millions cells were cross-linked by 4% formaldehyde solution at 37°C for 10 minutes followed by quenching with 125mM glycine. Cells were washed with PBS twice and harvested with scraper on ice. Cross-linked cells were resuspended in protease inhibitor containing SDS lysis buffer and subjected to sonication on a Diagenode bioruptor for 30 cycles at 30 seconds on followed by 30 seconds off. Sonication was standardized to yield fragments with mean length 150±50 bps. Sonicated lysate was cleared by centrifugation at 16000 rcf for 5 minutes at 4°C. 33 ul of lysate was reserved from each sample for input. CTCF ChIP was performed by using 150 µl of sonicated lysate. Sonicated lysate was incubated overnight at 4°C with antibody-conjugated beads. Beads were washed thrice with IP wash buffer 1 followed by washing with IP wash buffer 2 two times. Cross-links were reversed in reverse-crosslinking buffer for 15 mins at 55C followed by Proteinase K digestion at 65C for 15 mins. Reverse cross linked DNA was purified by DNA purification magnetic beads and used for library preparation and sequencing.

### Histone ChIP-sequencing

HEK293T cells transduced with control shRNA lentivirus, CGGBP1-shRNA lentivirus were selected with puromycin (10 µg/ml). Approximately 100 millions cells were cross-linked by 4% formaldehyde solution at 37°C for 10 minutes followed by quenching with 125mM glycine. Cells were washed with PBS twice and harvested with scraper on ice. Cross-linked cells were resuspended in protease inhibitor containing lysis buffer and subjected to sonication on a Diagenode bioruptor for 30 cycles at 30 seconds on followed by 30 seconds off to obtain mean sonicated chromatin of length 150±50 bps. Sonicated lysate was cleared by centrifugation at 16000 rcf for 5 minutes at 4°C. Sonicated lysates was pre-cleared by incubating with non-specific antibody/IgG and protein G sepharose beads. The pre-cleared lysate was incubated overnight at 4°C with antibody conjugated beads. Beads were washed with following buffers: low-salt IP wash buffer (0.1% SDS,1% Triton X-100, 2mM EDTA, 20mM Tris-HCl and 150mM NaCl), high-salt IP wash buffer (0.1% SDS,1% Triton X-100, 2mM EDTA, 20mM Tris-HCl and 500mM NaCl), LiCl IP wash buffer (0.25M LiCl, 1% IGEPAL, 1% Na deoxycholate, 1mM EDTA, 10mM Tris-HCl). Beads were washed with TE buffer twice (10mM Tris-HCl and 1mM EDTA). Beads-bound DNA was eluted in elution buffer (1% SDS and 0.1 M NaHCO_3_). Cross-links were reversed and followed by Proteinase K digestion. Reverse cross linked DNA was purified by DNA purification kit (Promega A1460) and used for library preparation and sequencing.

### Ion Torrent S5 library preparation and sequencing

The Ion XpressTM Plus Fragment Library Kit (Cat. no. 4471269) was used for library preparation from the above mentioned ChIP DNA samples. All the steps were performed as per manufacturer’s instructions for sequencing on Ion Torrent S5 platform without any barcoding. Briefly, the DNA were subjected to end-repair. The purified end-repaired DNA was ligated to Ion Torrent platform-compatible adapters. Nick repair was carried out to ensure the linking of the barcode adapters and DNA inserts on both the strands. The library was then amplified by PCR (number of cycles restricted to less than 18). AMPure XP beads were used to purify the PCR products (two rounds). The size-selected fragments were used for downstream clonal amplification on Ion Sphere Particles in an emulsion PCR. Prepared samples were sequenced on the Ion Proton S5 sequencer.

### Quality control and mapping of the ChIP-sequencing reads

Unpaired sequencing reads were trimmed and subjected to proprietary quality filtration for the Ion Torrent platform (minimum quality threshold 20). Reads with sequence length ranging from 80 to 300 base were used for further analysis. Reads were mapped using bowtie2 against the repeat-masked (hg38.fa.masked) or repeat-unmasked (hg38.fa) human genome as described. Reads were mapped end-to-end with default options.

### Analyses of genomic coordinates and sequences of regions

Sequences of bed coordinates were extracted from Hg38 (repeat-masked or unmasked as described) using bedtools getfasta tool. To isolate exclusive and overlapping CTCF peaks for datasets, bedtools intersect tool was used. To identify the number of overlapping ChIP-seq reads in any bed coordinate, bedtools coverage tool was used with fraction of overlap option. The closest distance between two different sets of bed coordinates were obtained using bedtools closest tool.

### Repeat content analyses

The repeat-masked and unmasked genome hg38 were used from UCSC genome browser. Sequences were repeat-masked using locally installed version of RepeatMasker. The repeat search engine used was RMBLAST (NCBI) and the repeat database used was obtained from RepBase.

### Motif finding

*De-novo* motif search was done using locally installed versions of MEME (version 5.0.3) suite tools meme and dreme. Suite tool fimo was used to find predicted motifs in sequence datasets. The motif search was performed using default options with -k value ranges from 12 to 15. Suite tool Tomtom was used to search for motif-corresponding transcription factors at HoCoMoCo database for transcription factor motifs.

### Plotting of signals in genomic coordinates

CTCF and histone modification reads signals were plotted along genomic coordinates using deepTools. Bam to BigWig conversions were done using bamcoverage tool. Matrices were generated by using computeMatrix in deepTools. The plots on these matrices were generated by using plotHeatmap (for heatmap) and plotProfile (average type summary plot) functions.

### Statistics and Graphs

Statistical tests were performed using Prism 8 (GraphPad) on numerical data generated from the above mentioned tools including OpenOffice Spreadsheet. R-scripts were used to plot distribution of CTCF reads in 0.5 kb bins from 1 million bins randomly selected from the whole genome (ggplot2), PCA analysis and hierarchical clustering (DiffBind) and correlation analysis of CTCF ChIP-seq datasets. Visualization of genomic features was carried out by the GUI version of locally installed Integrated Genome Viewer.

### Antibodies

Antibodies used in this study are as follows: CGGBP1 western blot (Proteintech 10716-1-AP), CGGBP1 IP (Proteintech 10716-1-AP; Santacruz SC-376482), CTCF western blot (SC-271514), CTCF IP (Santa Cruz SC-28198 and SC-271514), CTCF ChIP (Santa Cruz SC-28198 and SC-271514), H3K4me3 ChIP (ABCAM ab8580), H3K9me3 ChIP (SantaCruz SC-130356) and H3K27me3 ChIP (ABCAM ab6002), IgG ChIP preincubation (FBS from Himedia), GAPDH (NB300-328SS), Mouse and Rabbit IgG in PLA (Invitrogen).

### Publicly available data usage

The following publicly available datasets were used in this study: ChIP-seq datasets from ENCODE (ENCFF567GON, ENCFF757KYL, ENCFF510QXG, ENCFF712LFQ, ENCFF687IUD, ENCFF420KMT, ENCFF217ZMF, ENCFF180BYN, ENCFF987YIJ, ENCFF002CVC, ENCFF351VGZ, ENCFF002CVE, ENCFF082IQD, ENCFF474PPT, ENCFF002CVF, ENCFF560WFS, ENCFF895JAW, ENCFF139EBY, ENCFF380ZXB and ENCFF938BOJ), UCSC CTCF Binding Sites (https://genome-test.gi.ucsc.edu/cgi-bin/hgTables?command=start), UCSC Regulatory Elements (available through UCSC table Browser). Human genome hg38 was downloaded from UCSC. UCSC LiftOver tool was used to convert genomic coordinates between different versions of the human genome. CGGBP1 ChIP-seq datasets were obtained from NCBI Geo Datasets (GSE53571).

