## Supplementary tables and legends for "CGGBP1 regulates chromatin barrier activity and CTCF occupancy at repeats"

| Sample Name | Total Reads | Mapped Reads | Unmapped Reads | Percentage of mapped reads | Percentage of unmapped reads |
|---|---|---|---|---|---|
| **RM CTCF CT** | 99086307 | 33436004 | 65650303 | 33.74 | 66.25 |
| **RM CTCF KD** | 76777977 | 31636108 | 45141869 | 41.20 | 58.79 |
| **RM CTCF OE** | 74091715 | 28146793 | 45944922 | 37.98 | 62.01 |

Table S1: The table shows number and percentage of mapped and unmapped CTCF reads to repeat-masked human genome (hg38).

| Sample Name | Number of RM CTCF peaks | Number of CTCF motifs in CTCF peak sequences | Average number of motifs/peak |
|---|---|---|---|
| **RM CTCF CT** | 26635 | 15187 | 0.57 |
| **RM CTCF KD** | 24418 | 23563 | 0.96 |
| **RM CTCF OE** | 21071 | 18983 | 0.90 |

Table S2: Repeat-masked (RM) CTCF reads were used to call peaks. The table shows number of repeat-masked CTCF peaks and CTCF motifs count in peak sequences.

| Sample Name | Number of RM CTCF peaks | Number of L1 motifs in peaks | Motif positive peaks | Percentage peaks with Alu-L1 motifs | Average number of motifs/peak |
|---|---|---|---|---|---|
| **RM CTCF CT** | 26635 | 18986 | 4413 | 16.56 | 4.30 |
| **RM CTCF KD** | 24418 | 36253 | 13787 | 56.46 | 2.63 |
| **RM CTCF OE** | 21071 | 34391 | 11550 | 54.81 | 2.98 |

Table S3: Table shows number of L1-motifs and percentage of peaks with L1 motifs for repeat-masked CTCF peaks for CT, KD and OE.

| Sample Name | Total Reads | Mapped Reads | Unmapped Reads | Percentage of mapped reads | Percentage of unmapped reads |
|---|---|---|---|---|---|
| **CTCF CT** | 99086307 | 79778533 | 19307774 | 80.51 | 19.49 |
| **CTCF KD** | 76777977 | 74389175 | 2388802 | 96.89 | 3.11 |
| **CTCF OE** | 74091715 | 67600036 | 6491679 | 91.24 | 8.76 |
| **RM CTCF CT** | 99086307 | 33436004 | 65650303 | 33.74 | 66.25 |
| **RM CTCF KD** | 76777977 | 31636108 | 45141869 | 41.20 | 58.79 |
| **RM CTCF OE** | 74091715 | 28146793 | 45944922 | 37.98 | 62.01 |

Table S4: The table shows number and percentage of mapped and unmapped CTCF reads to repeat-unmasked and repeat-masked (RM) human genome (hg38)

| | CTCF CT | CTCF KD | CTCF OE |
|---|---|---|---|
| **SINEs** | 6.1 | 3.53 | 3.11 |
| ALUs | 4.85 | 2.19 | 1.64 |
| MIRs | 1.23 | 1.3 | 1.43 |
| **LINEs** | 19.14 | 9.65 | 10.46 |
| LINE1 | 17.79 | 8.32 | 8.99 |
| LINE2 | 1.17 | 1.15 | 1.27 |
| L3/CR1 | 0.13 | 0.14 | 0.14 |
| **LTR elements** | 4.65 | 4.99 | 5.1 |
| ERVL | 1.04 | 1.4 | 1.41 |
| ERVL-MaLRs | 1.33 | 1.1 | 1.36 |
| ERV_classI | 1.46 | 1.48 | 1.42 |
| ERV_classII | 0.68 | 0.82 | 0.73 |
| **DNA elements** | 2.28 | 2.4 | 2.35 |
| hAT-Charlie | 1.14 | 1.33 | 1.24 |
| TcMar-Tigger | 0.61 | 0.55 | 0.61 |
| **Unclassified** | 0.38 | 0.1 | 0.09 |
| **Total interspersed repeats** | 32.54 | 20.66 | 21.11 |
| **Small RNA** | 0.16 | 0.22 | 0.24 |
| **Satellites** | 11.48 | 11.49 | 14.4 |
| **Simple repeats** | 0 | 0 | 0 |
| **Low complexity** | 0 | 0 | 0 |

Table S5: Repeat content in CTCF peaks for CT, KD and OE samples.

| Sample Name | Number of CTCF peaks | Number of CTCF motifs in CTCF peak sequences |
|---|---|---|
| **CTCF CT** | 42978 | 12807 |
| **CTCF KD** | 47632 | 36405 |
| **CTCF OE** | 47216 | 21767 |

Table S6:  Repeat-unmasked (RM) CTCF reads were used to call peaks. The table shows number of CTCF peaks and CTCF motifs  count in peak sequences.

| | CTvsKD peaks with motifs | CTvsOE peaks with motifs | KDvsCT peaks with motifs | KDvsOE peaks with motifs | OEvsCT peaks with motifs | OEvsKD peaks with motifs |
|---|---|---|---|---|---|---|
| **SINEs** | 8.76 | 6.52 | 3.36 | 3.65 | 2.77 | 3.28 |
| ALUs | 7.12 | 5.28 | 1.83 | 2.16 | 1.34 | 1.29 |
| MIRs | 1.59 | 1.21 | 1.47 | 1.43 | 1.38 | 1.93 |
| **LINEs** | 8.08 | 7.02 | 5.37 | 5.96 | 5.29 | 8.23 |
| LINE1 | 6.54 | 5.61 | 3.87 | 4.43 | 3.81 | 6.07 |
| LINE2 | 1.32 | 1.26 | 1.33 | 1.35 | 1.27 | 1.69 |
| L3/CR1 | 0.22 | 0.14 | 0.14 | 0.16 | 0.16 | 0.32 |
| **LTR elements** | 7.93 | 7.93 | 7.24 | 8.66 | 6.57 | 8.53 |
| ERVL | 1.6 | 1.78 | 1.72 | 1.96 | 1.81 | 2.25 |
| ERVL-MaLRs | 2.51 | 2.02 | 1.81 | 2.14 | 1.67 | 2.61 |
| ERV_classI | 2.61 | 2.31 | 2.18 | 2.73 | 1.82 | 2.62 |
| ERV_classII | 1.14 | 1.6 | 1.26 | 1.51 | 0.97 | 0.76 |
| **DNA elements** | 2.47 | 2.37 | 2.64 | 2.54 | 3.02 | 3.11 |
| hAT-Charlie | 1.68 | 1.53 | 1.47 | 1.34 | 1.86 | 1.99 |
| TcMar-Tigger | 0.38 | 0.37 | 0.55 | 0.59 | 0.51 | 0.42 |
| **Unclassified** | 0.07 | 0.07 | 0.08 | 0.11 | 0.06 | 0.07 |
| **Total interspersed repeats** | 27.3 | 23.91 | 18.7 | 20.92 | 17.72 | 23.22 |
| **Small RNA** | 0.11 | 0.1 | 0.04 | 0.04 | 0.05 | 0.04 |
| **Satellites** | 12.3 | 10.97 | 2.95 | 5.64 | 2.28 | 6.98 |
| **Simple repeats** | 0 | 0 | 0 | 0 | 0 | 0 |
| **Low complexity** | 0 | 0 | 0 | 0 | 0 | 0 |

Table S7: Repeat content in  exclusive CTCF peaks with CTCF motif for CT, KD and OE.

| | CTvsKD peaks no motifs | CTvsOE peaks no motifs | KDvsCT peaks no motifs | KDvsOE peaks no motifs | OEvsCT peaks no motifs | OEvsKD peaks no motifs |
|---|---|---|---|---|---|---|
| **SINEs** | 9.24 | 9.18 | 4.55 | 4.97 | 4.05 | 4.25 |
| ALUs | 7.74 | 7.71 | 3.05 | 3.57 | 2.14 | 2.27 |
| MIRs | 1.46 | 1.45 | 1.46 | 1.38 | 1.89 | 1.96 |
| **LINEs** | 30.95 | 30.67 | 14.77 | 19.75 | 15.63 | 19.5 |
| LINE1 | 29.2 | 28.95 | 13.04 | 17.98 | 13.56 | 17.33 |
| LINE2 | 1.5 | 1.49 | 1.44 | 1.47 | 1.78 | 1.85 |
| L3/CR1 | 0.17 | 0.17 | 0.22 | 0.23 | 0.2 | 0.22 |
| **LTR elements** | 5.27 | 5.25 | 5.15 | 5.1 | 6.12 | 6.11 |
| ERVL | 0.83 | 0.84 | 1.53 | 1.27 | 1.48 | 1.28 |
| ERVL-MaLRs | 2.04 | 2.01 | 1.42 | 1.52 | 2.2 | 2.39 |
| ERV_classI | 1.83 | 1.83 | 1.67 | 1.73 | 1.83 | 1.85 |
| ERV_classII | 0.44 | 0.43 | 0.4 | 0.48 | 0.45 | 0.45 |
| **DNA elements** | 2.75 | 2.73 | 3.05 | 2.92 | 2.74 | 2.8 |
| hAT-Charlie | 1.2 | 1.18 | 1.6 | 1.43 | 1.18 | 1.17 |
| TcMar-Tigger | 0.93 | 0.9 | 0.88 | 0.93 | 1.04 | 1.15 |
| **Unclassified** | 0.67 | 0.69 | 0.1 | 0.15 | 0.1 | 0.12 |
| **Total interspersed repeats** | 48.87 | 48.52 | 27.62 | 32.9 | 28.64 | 32.78 |
| **Small RNA** | 0.05 | 0.03 | 0.21 | 0.07 | 0.2 | 0.09 |
| **Satellites** | 7.89 | 7.84 | 13.46 | 12.65 | 18.92 | 18.61 |
| **Simple repeats** | 0 | 0 | 0 | 0 | 0 | 0 |
| **Low complexity** | 0 | 0 | 0 | 0 | 0 | 0 |

Table S8: Repeat content in exclusive CTCF peaks without CTCF motif for CT, KD and OE.

| Sample Name | Number of CTCF peaks | Number of L1 motifs in peaks | Motif positive peaks | Percentage peaks with Alu-L1 motifs | Average number of motifs/peak |
|---|---|---|---|---|---|
| **CTCF CT** | 42978 | 96178 | 28798 | 67.01 | 3.34 |
| **CTCF KD** | 47632 | 93417 | 31525 | 66.19 | 2.96 |
| **CTCF OE** | 47216 | 103395 | 30450 | 64.49 | 3.40 |

Table S9: Table shows number of L1-motifs and percentage of peaks with L1 motifs for CTCF peaks for CT, KD and OE.

| Sample Name | Total Reads | Mapped Reads | Unmapped Reads | Percentage of mapped reads | Percentage of unmapped reads |
|---|---|---|---|---|---|
| **H3K4me3 CT** | 110040103 | 100149169 | 9890934 | 91.01151877 | 8.988481227 |
| **H3K4me3 KD** | 93091534 | 90279250 | 2812284 | 96.97901208 | 3.020987924 |
| **H3K9me3 CT** | 82117627 | 69952262 | 12165365 | 85.18544015 | 14.81455985 |
| **H3K9me3 KD** | 80525298 | 61677565 | 18847733 | 76.59402266 | 23.40597734 |
| **H3K27me3 CT** | 95003513 | 83135564 | 11867962 | 87.50787037 | 12.49212963 |
| **H3K27me3 KD** | 66148370 | 63752690 | 2395680 | 96.37832346 | 3.621676543 |

Table S10: The table shows number and percentage of mapped and unmapped histone modification ChIP-seq reads to repeat-unmasked human genome (hg38).

| | Loss of CTCF binding | Gain in CTCF binding |
|---|---|---|
| **SINEs** | 9.54 | 2.34 |
| ALUs | 8.02 | 1.54 |
| MIRs | 1.52 | 0.81 |
| **LINEs** | 35.3 | 10.47 |
| LINE1 | 34.38 | 9.36 |
| LINE2 | 0.78 | 1.11 |
| L3/CR1 | 0.09 | 0 |
| **LTR elements** | 5.9 | 6.81 |
| ERVL | 0.81 | 0.95 |
| ERVL-MaLRs | 1.8 | 1.7 |
| ERV_classI | 2.39 | 2.52 |
| ERV_classII | 0.78 | 1.31 |
| **DNA elements** | 2.9 | 1.77 |
| hAT-Charlie | 1.48 | 1.29 |
| TcMar-Tigger | 0.86 | 0.31 |
| **Unclassified** | 0.44 | 0 |
| **Total interspersed repeats** | 54.08 | 21.4 |
| **Small RNA** | 0.04 | 0 |
| **Satellites** | 1.35 | 4.75 |
| **Simple repeats** | 0 | 0 |
| **Low complexity** | 0 | 0 |

Table S11: Repeat content in exclusive CTCF peaks with differential H3K9me3 signal in flanks undergoing gain or loss of boundary element function upon CGGBP1 knockdown.
