## Supplementary figures and legends for "CGGBP1 regulates chromatin barrier activity and CTCF occupancy at repeats"

#### Figure S1:

A

B

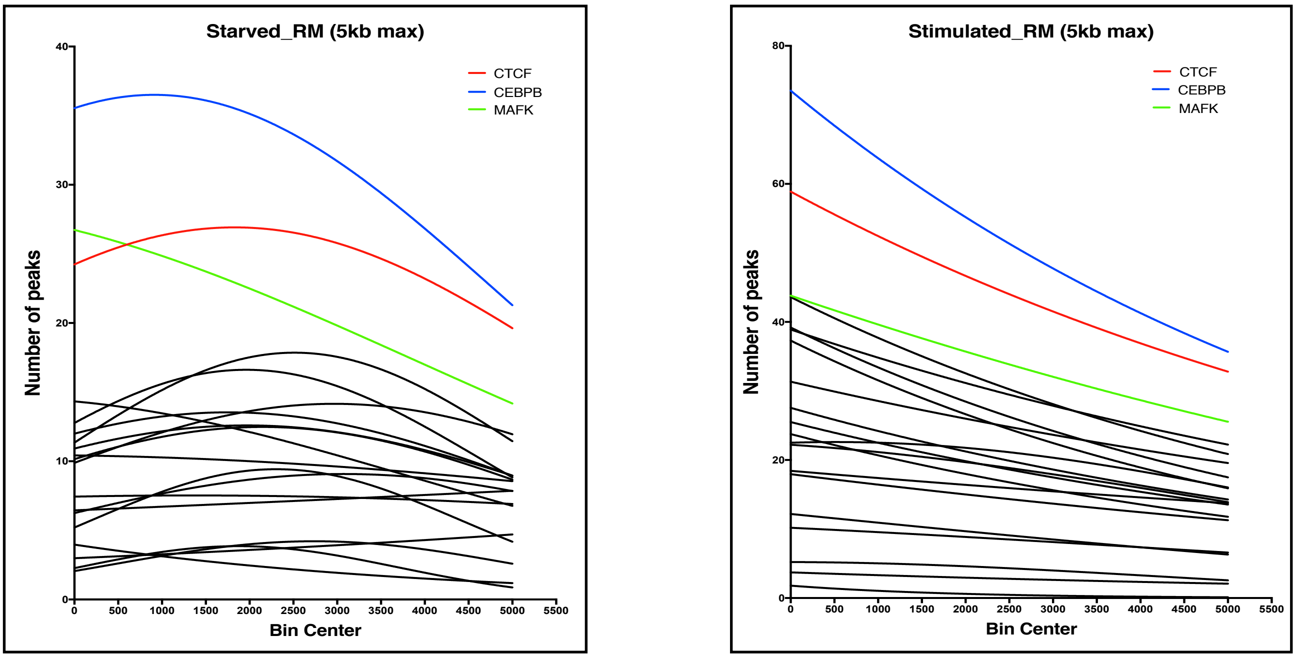


C

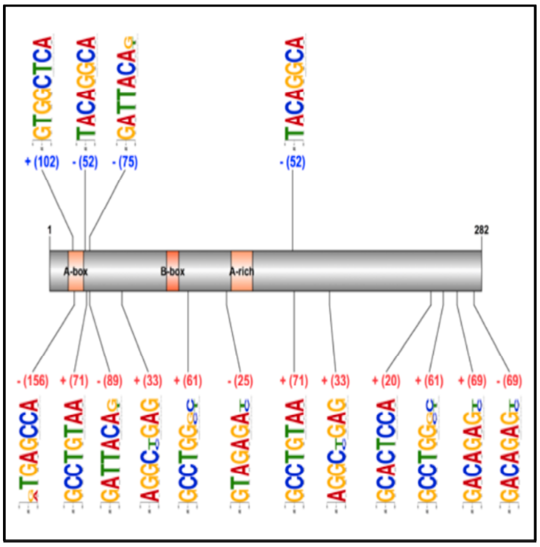


Number of Alu subsequences (SS) = 8

% of RM stimulated CGGBP1 peaks with Alu SS = 20.02

% of RM CT peaks with Alu SS = 6.83

% of RM KD peaks with Alu SS = 23.20

% of RM OE peaks with Alu SS = 22.03

D

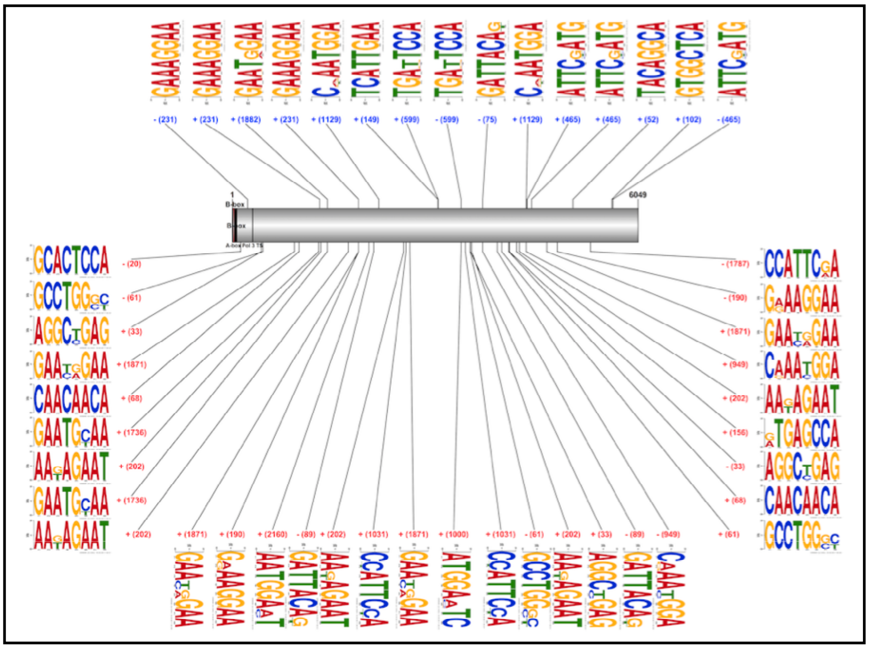


Number of L1 subsequences (SS) = 15

% of RM stimulated CGGBP1 peaks with L1 SS = 44.38

% of RM CT peaks with L1 SS = 15.48

% of RM KD peaks with L1 SS = 52.67

% of RMOE peaks with L1 SS = 51.03

#### Figure S2:

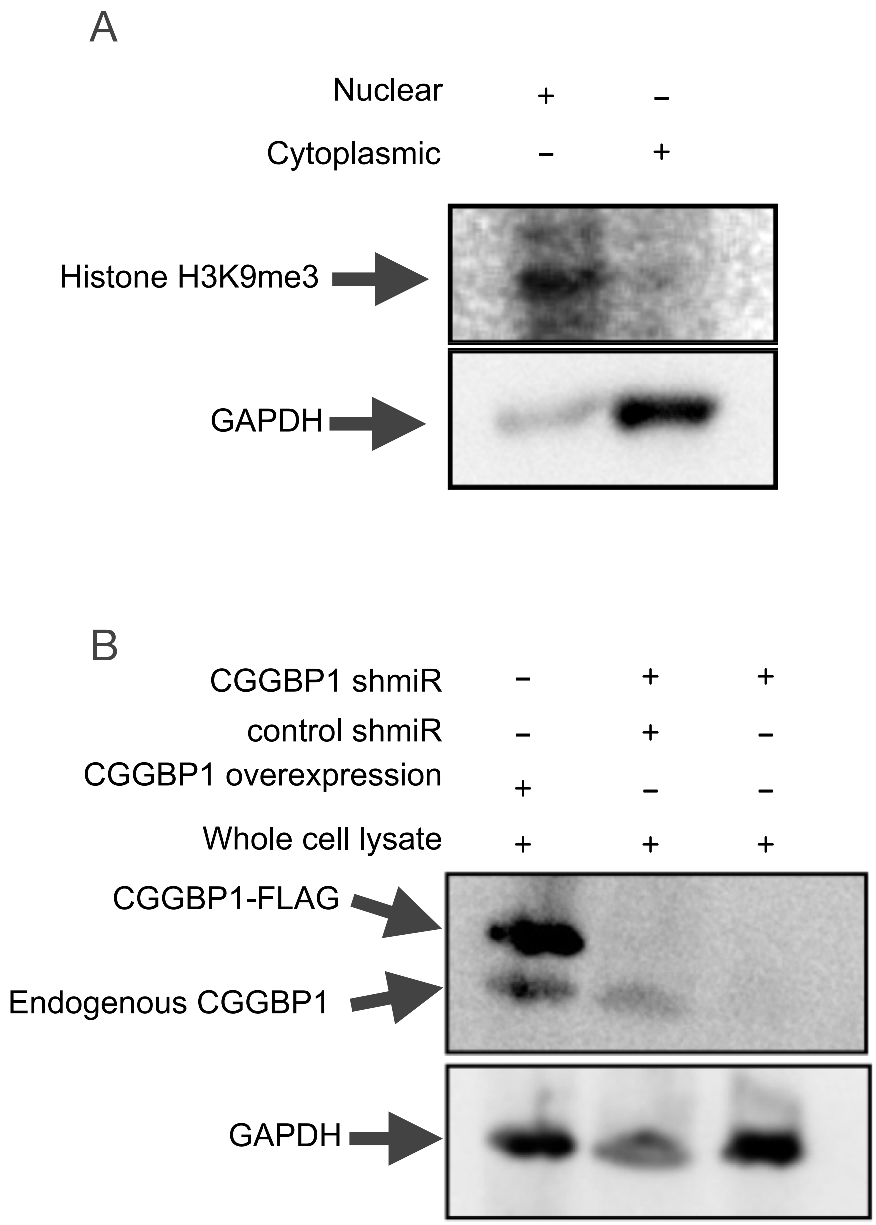
A
Nuclear + –
Cytoplasmic – +
Histone H3K9me3
GAPDH
B
CGGBP1 shmiR – + +
control shmiR – + –
CGGBP1 overexpression + – –
Whole cell lysate + + +
CGGBP1-FLAG
Endogenous CGGBP1
GAPDH

**Figure S3:**

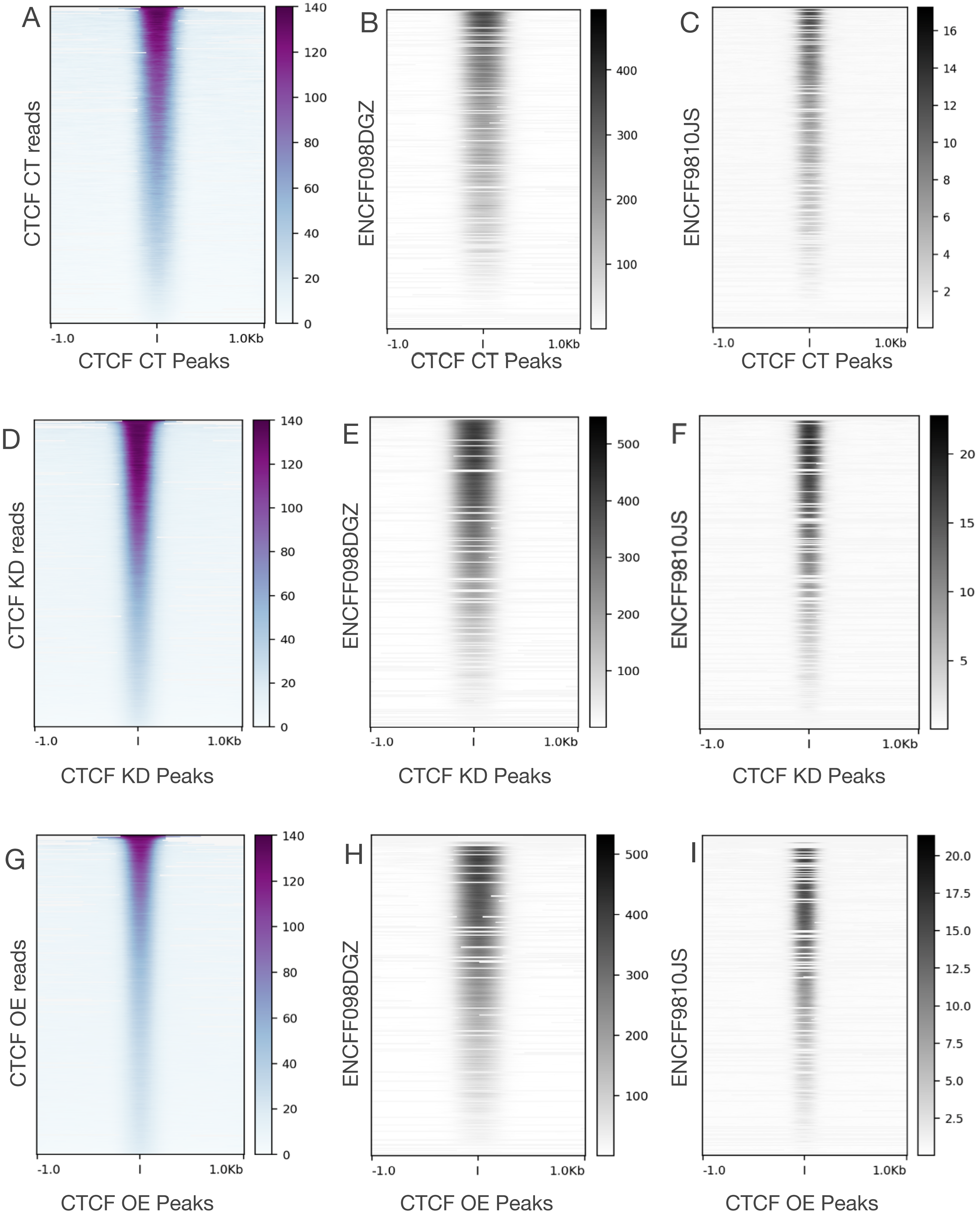

A
CTCF CT reads
CTCF CT Peaks
B
ENCFF098DGZ
CTCF CT Peaks
C
ENCFF9810JS
CTCF CT Peaks
D
CTCF KD reads
CTCF KD Peaks
E
ENCFF098DGZ
CTCF KD Peaks
F
ENCFF9810JS
CTCF KD Peaks
G
CTCF OE reads
CTCF OE Peaks
H
ENCFF098DGZ
CTCF OE Peaks
I
ENCFF9810JS
CTCF OE Peaks
-1.0
1.0Kb

#### Figure S4:

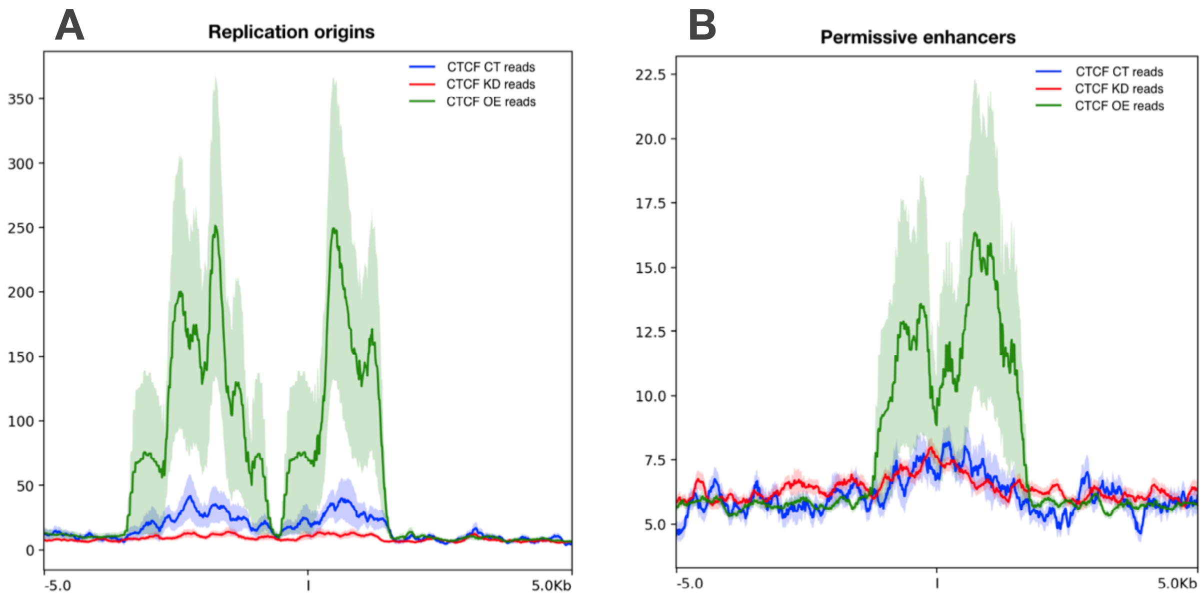

A
Replication origins
CTCF CT reads
CTCF KD reads
CTCF OE reads
350
300
250
200
150
100
50
0
-5.0
5.0Kb
B
Permissive enhancers
CTCF CT reads
CTCF KD reads
CTCF OE reads
22.5
20.0
17.5
15.0
12.5
10.0
7.5
5.0
-5.0
5.0Kb

### Figure S5:

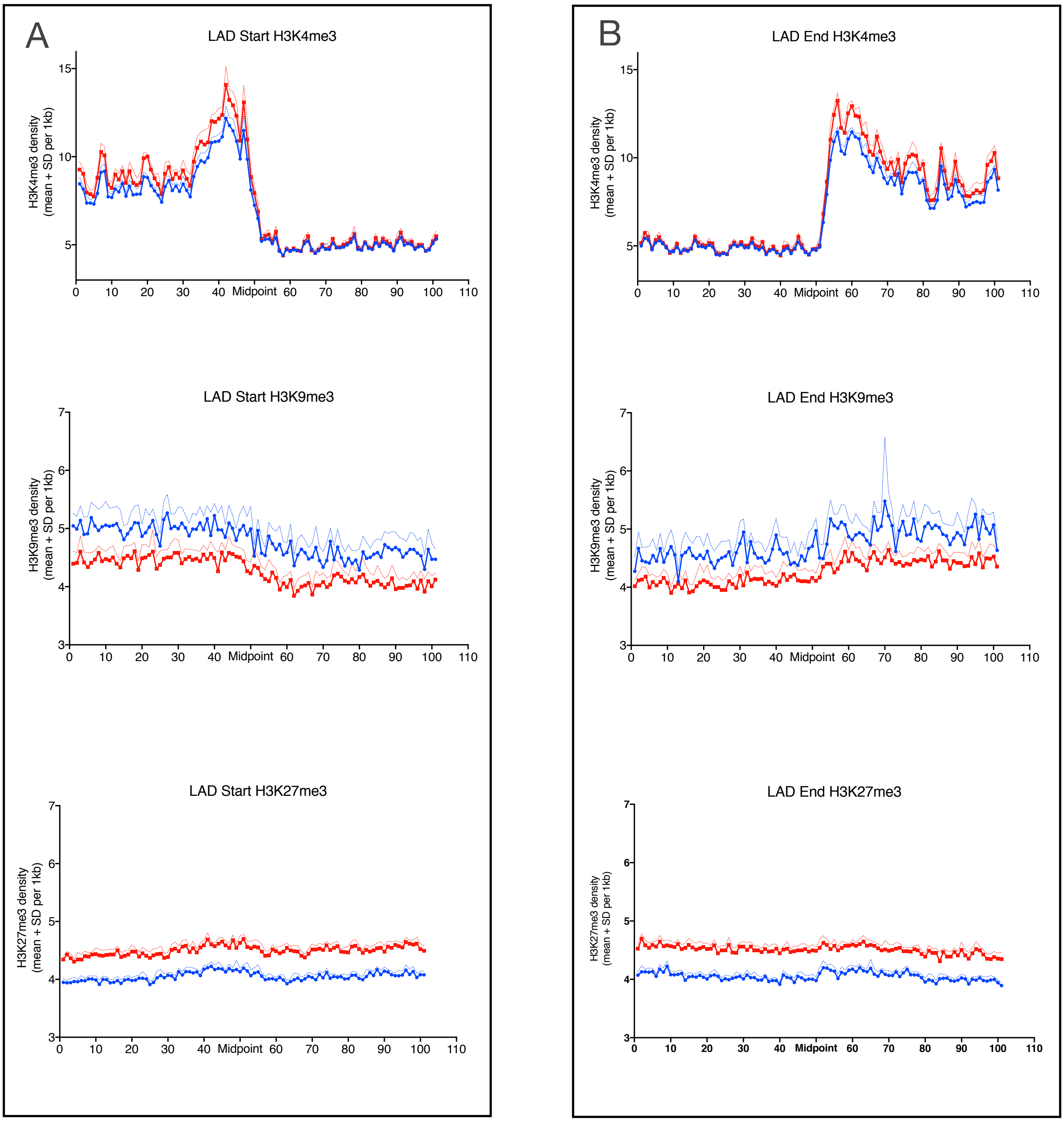

A
LAD Start H3K4me3
H3K4me3 density (mean + SD per 1kb)
0 10 20 30 40 Midpoint 60 70 80 90 100 110
LAD Start H3K9me3
H3K9me3 density (mean + SD per 1kb)
0 10 20 30 40 Midpoint 60 70 80 90 100 110
LAD Start H3K27me3
H3K27me3 density (mean + SD per 1kb)
0 10 20 30 40 Midpoint 60 70 80 90 100 110
B
LAD End H3K4me3
H3K4me3 density (mean + SD per 1kb)
0 10 20 30 40 Midpoint 60 70 80 90 100 110
LAD End H3K9me3
H3K9me3 density (mean + SD per 1kb)
0 10 20 30 40 Midpoint 60 70 80 90 100 110
LAD End H3K27me3
H3K27me3 density (mean + SD per 1kb)
0 10 20 30 40 Midpoint 60 70 80 90 100 110

**Figure S6:**

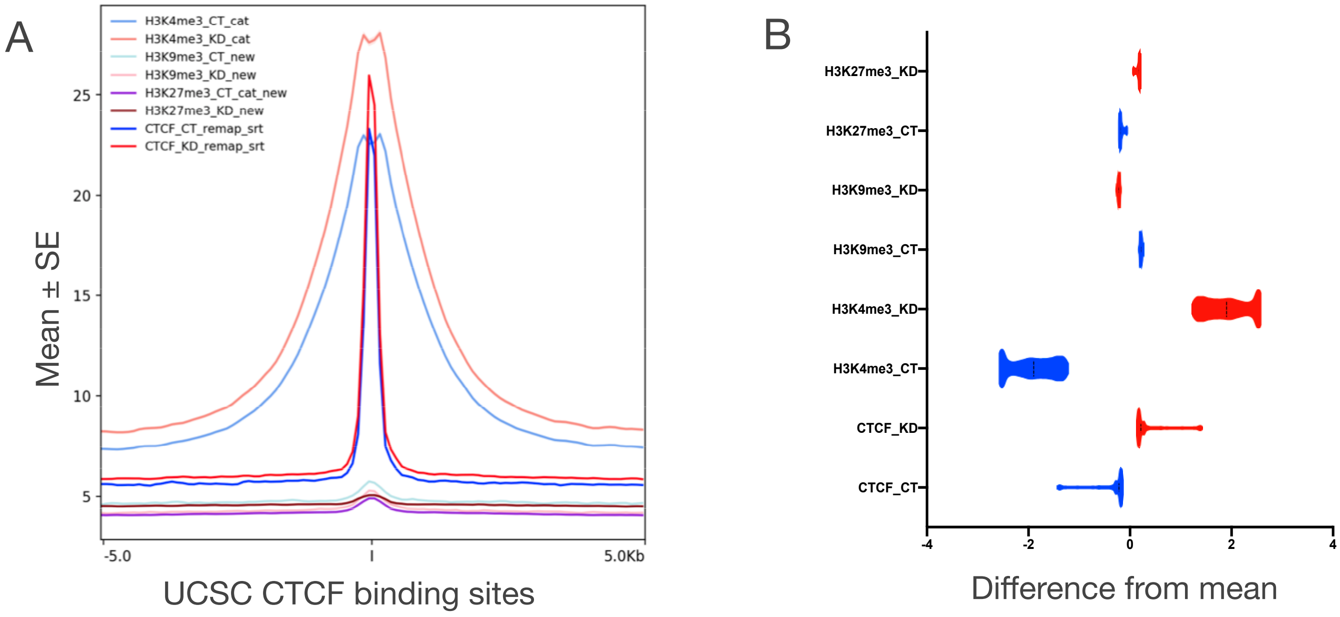
A
H3K4me3_CT_cat
H3K4me3_KD_cat
H3K9me3_CT_new
H3K9me3_KD_new
H3K27me3_CT_cat_new
H3K27me3_KD_new
CTCF_CT_remap_srt
CTCF_KD_remap_srt
25
20
15
10
5
Mean ± SE
-5.0
5.0Kb
UCSC CTCF binding sites
B
H3K27me3_KD
H3K27me3_CT
H3K9me3_KD
H3K9me3_CT
H3K4me3_KD
H3K4me3_CT
CTCF_KD
CTCF_CT
-4
-2
0
2
4
Difference from mean

#### Figure S7:

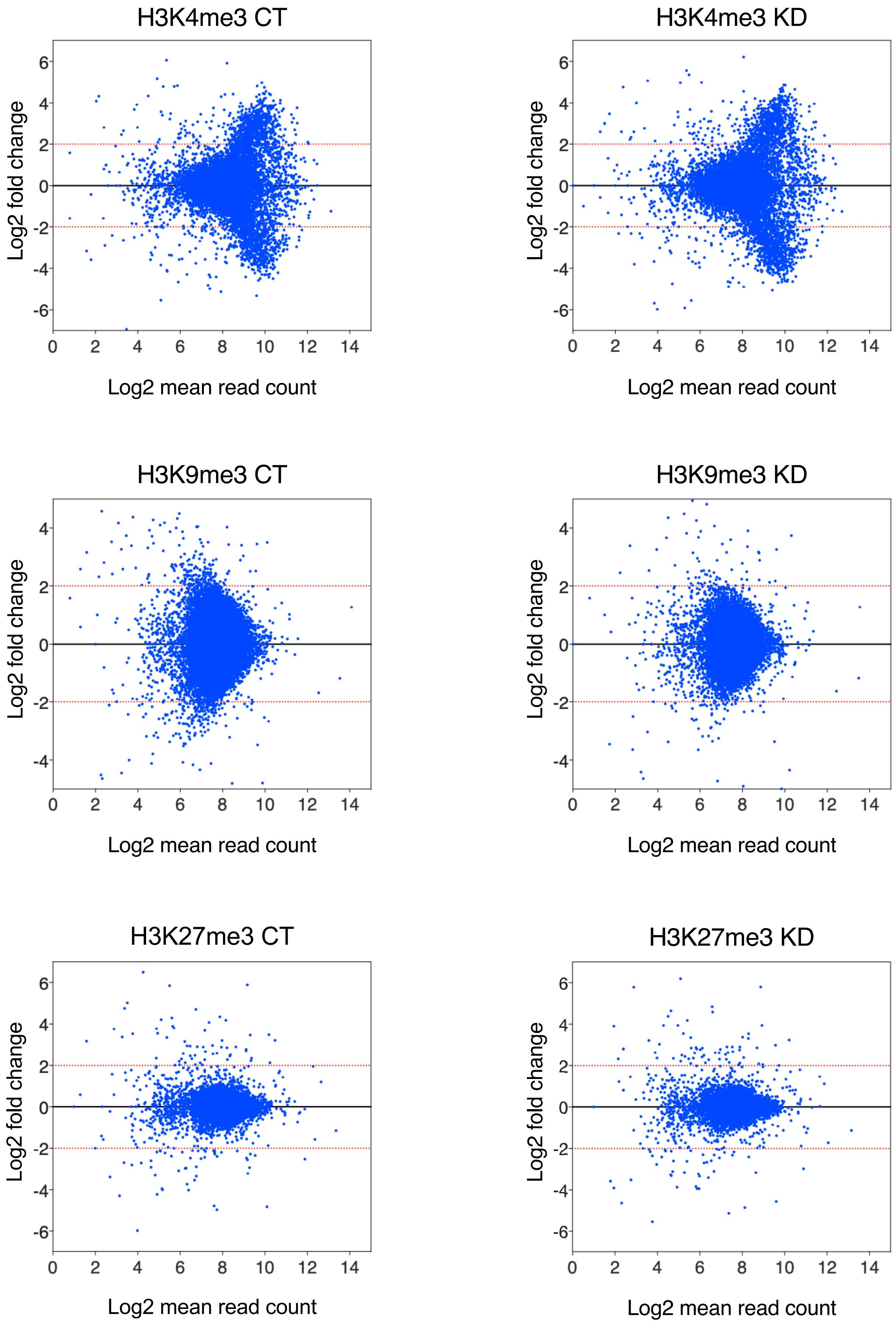
H3K4me3 CT
H3K4me3 KD
H3K9me3 CT
H3K9me3 KD
H3K27me3 CT
H3K27me3 KD
Log2 fold change
Log2 mean read count
0
2
4
6
8
10
12
14
6
4
2
0
-2
-4
-6

### Figure S8:

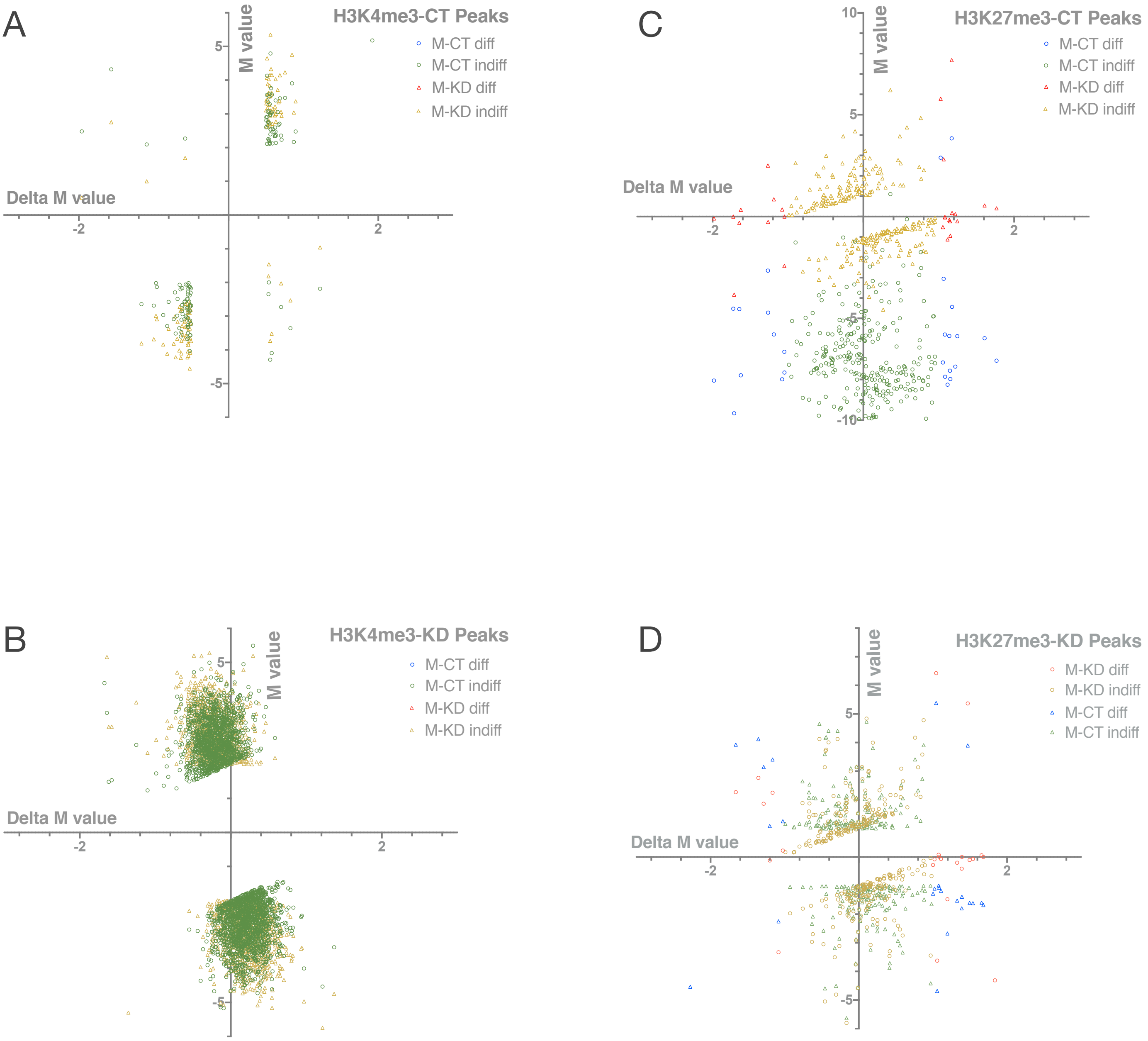
A
H3K4me3-CT Peaks
M-CT diff
M-CT indiff
M-KD diff
M-KD indiff
Delta M value
M value
B
H3K4me3-KD Peaks
M-CT diff
M-CT indiff
M-KD diff
M-KD indiff
Delta M value
M value
C
H3K27me3-CT Peaks
M-CT diff
M-CT indiff
M-KD diff
M-KD indiff
Delta M value
M value
D
H3K27me3-KD Peaks
M-KD diff
M-KD indiff
M-CT diff
M-CT indiff
Delta M value
M value

Supplementary Figure Legends

Figure S1: (A and B) The closest distance between Starved RM CGGBP1 peak midpoint and transcription factor peak midpoint was determined by bedtools closest. Frequency distribution of closest distances was plotted in bin of 0.5 for starved RM CGGBP1 peaks (A) and stimulated RM CGGBP1 peaks (B). (C and D) RM-CGGBP1 peaks with tag count more than 10 were were splitted into narrow and broad peak based on peak-summit location. Narrow peak have peak-summit in central ⅓ region of peak length (y/x = 1.5 to 2.5). Rest of the peaks were annotated as broad peaks. Central coordinates sequences of narrow peaks (Peak start + 0.4 peak length to Peak-start +0.667 peak-length) were fetched from RM genomes. RM CGGBP1 narrow peak-central coordinates sequences were subjected to De-novo motif search by using DREME (minK 8 ). RM-CGGBP1 narrow peaks in starved and stimulated condition were centrally enriched with repeat-derived DNA motifs which aligned to consensus sequences of Alu-SINE (C) and L1-LINE (D).

Figure S2: (A) Cytoplasmic and nuclear fraction were separated from HEK293T cells by REAP protocol. The upper panel shows immunoblot results for nuclear marker H3K9me3 and lower panel shows immunoblot results for cytoplasmic marker GAPDH. (B) HEK293 cells were transduced with control shRNA lentivirus, CGGBP1-shRNA lentivirus and CGGBP1 over-expression lentivirus respectively. The upper panel shows immunoblot results for CGGBP1 and lower panel shows same for GAPDH loading control.

Figure S3: (A to C) Distribution of CTCF CT reads at unmasked CTCF CT-peaks was plotted for 1kb flanks in bin size of 10 (A). Similarly distribution of published CTCF reads in prostate epithelial cells (ENCFF098DGZ) (B) and A549 (ENCFF9810JS) (C) was also plotted at CTCF CT peaks. (D to F) Distribution of CTCF KD reads at unmasked CTCF KD-peaks was plotted for 1kb flanks in bin size of 10 (D). Similarly distribution of published CTCF reads in prostate epithelial cells (ENCFF098DGZ) (E) and A549 (ENCFF9810JS) (F) was also plotted at CTCF KD peaks. (G to I) Distribution of CTCF OE reads at unmasked CTCF OE-peaks was plotted for 1kb flanks in bin size of 10 (G). Similarly distribution of published CTCF reads in prostate epithelial cells (ENCFF098DGZ) (H) and A549 (ENCFF9810JS) (I) was also plotted at CTCF OE peaks.

Figure S4: Distribution of CTCF reads at replication origin was plotted for 5kb flanks in bin size of 10 (A). Distribution of CTCF reads at enhancers (UCSC Regulation datasets) was plotted for 5kb flanks in bin size of 10 (B).

Figure S5: (A and B) Distribution of Histone modification reads for CT and KD samples was plotted in 1 Mb upstream and downstream of  LAD boundary. Histone modification reads counts in bin size of 1kb was plotted for CT (blue) and KD (red) peaks at LAD start site (A) and LAD end site (B).

Figure S6: Distribution of CTCF and histone modification reads at UCSC CTCF binding sites was  plotted for 5 kb flanks in bin size  of 10 (A). Variation of CTCF and histone reads in bin size 10 at UCSC CTCF binding sites was shown as difference from mean for CT and KD sample (B).

Figure S7: Difference in histone modifications read occupancy in upstream and downstream 10kb flanks of the all exclusive peaks were compared between CT and KD. Reads coverage count for 10kb upstream and downstream was converted to logarithmic scale (log base = 2). Log2 fold change (M = Upstream-Downstream) was plotted against average read count for CT (A) and KD (B).

Figure S8: (A  and B) As described above, delta M value for CT/KD exclusive peaks were calculated for the H3K4me4 reads. Those peaks with significantly different (delta M value <-2 to >+2 ) H3K4me3 profile in CT and KD are represented as blue for CT (M-CT diff) and red for KD (M-KD diff), whereas those peaks which H3k4me3 profile did not altered significantly in KD (delta M value ranges from -2 to +2) are highlighted as green for CT (M-CT indiff) and yellow for KD (M-KD indiff) (A). Similarly, delta M value was calculated for KD/CT exclusive peaks and those peaks which showed substantial changes (delta M value <-2 and >+2 ) in H3K4me3 profile in KD are represented as red for KD (M-KD diff) and blue for CT (M-CT diff). Peaks which H3K4me3 profile did not changed substantially by CGGBP1 depletion are shown as yellow for KD (M-KD indiff) and green for CT (M-CT indiff) (B). (C  and D) Similarly, delta M value for CT/KD exclusive peaks were calculated for the H3K27me4 reads. Those peaks with significantly different (delta M value <-1 to >+1 ) H3K27me3 profile in CT and KD are represented as blue for CT (M-CT diff) and red for KD (M-KD diff), whereas those peaks which H3k27me3 profile did not altered significantly in KD (delta M value ranges from -1 to +1) are highlighted as green for CT (M-CT indiff) and yellow for KD (M-KD indiff) (A). Similarly, delta M value was calculated for KD/CT exclusive peaks and those peaks which showed substantial changes (delta M value <-1 and >+1 ) in H3K27me3 profile in KD are represented as red for KD (M-KD diff) and blue for CT (M-CT diff). Peaks which H3K27me3 profile did not changed substantially by CGGBP1 depletion are shown as yellow for KD (M-KD indiff) and green for CT (M-CT indiff) (B).
